# Inferring parameters of pyramidal neuron excitability in mouse models of Alzheimer’s disease using biophysical modeling and deep learning

**DOI:** 10.1101/2023.04.18.537149

**Authors:** Soheil Saghafi, Timothy Rumbell, Viatcheslav Gurev, James Kozloski, Francesco Tamagnini, Kyle C. A. Wedgwood, Casey O. Diekman

## Abstract

Alzheimer’s disease (AD) is believed to occur when abnormal amounts of the proteins amyloid beta and tau aggregate in the brain, resulting in a progressive loss of neuronal function. Hippocampal neurons in transgenic mice with amyloidopathy or tauopathy exhibit altered intrinsic excitability properties. We introduce a novel parameter inference technique, deep hybrid modeling (DeepHM), that combines deep learning with biophysical modeling to map experimental data recorded from hippocampal CA1 neurons in transgenic AD mice and age-matched wildtype littermate controls to the parameter space of a conductance-based CA1 model. Although mechanistic modeling and machine learning methods are by themselves powerful tools for approximating biological systems and making accurate predictions from data, when used in isolation these approaches suffer from distinct shortcomings: model and parameter uncertainty limit mechanistic modeling, whereas machine learning methods disregard the underlying biophysical mechanisms. DeepHM addresses these shortcomings by using conditional generative adversarial networks (cGANs) to provide an inverse mapping of data to mechanistic models that identifies the distributions of mechanistic modeling parameters coherent to the data. Here, we demonstrate that DeepHM accurately infers parameter distributions of the conductance-based model and outperforms a Markov chain Monte Carlo method on several test cases using synthetic data. We then use DeepHM to estimate parameter distributions corresponding to the experimental data and infer which ion channels are altered in the Alzheimer’s mouse models compared to their wildtype controls at 12 and 24 months. We find that the conductances most disrupted by tauopathy, amyloidopathy, and aging are delayed rectifier potassium, transient sodium, and hyperpolarization-activated potassium, respectively.

## 1 Introduction

Although the underlying cause of Alzheimer’s disease (AD) remains poorly understood, it is believed to occur when abnormal amounts of the proteins amyloid beta and tau aggregate in the brain, forming extracellular plaques (amyloidopathy) and neurofibrillary tangles (tauopathy) that result in a progressive loss of neuronal function and dementia [1, 2]. In transgenic mice with amyloidopathy, neurons in the hippocampus—a brain structure critical for memory—exhibit altered intrinsic excitability properties, such as action potentials with reduced peaks and widths [3–5]. Hippocampal neurons in transgenic mice with tauopathy also show altered excitability, but in different properties such as hyperpolarization-activated membrane potential sag and action potential threshold [6, 7].

Ideally, biophysical modeling could be used to gain insights into the mechanisms underlying the disrupted electrophysiological properties of these Alzheimer’s mutant mice. However, determining whether or not such a biophysical model and its outputs are coherent with a set of experimental observations is a major challenge since such models contain many unknown parameters and are not amenable to statistical inference due to their non-invertibility. The main difficulty in solving the inverse problem for mechanistic models arises from intractability of the likelihood function [8]. On the other hand, neither purely statistical models with tractable likelihoods nor purely data-driven machine learning algorithms offer much insight into underlying biological mechanisms [9, 10]. Here, we use deep learning to perform inversion of complex biophysical models and enable the mapping of experimental data into the space of biophysical model parameters. Since this approach combines deep learning with mechanistic modeling, we refer to it as deep hybrid modeling (DeepHM).

In biological systems, the tremendous amount of inherent cell-to-cell variability presents a significant challenge to mapping experimental data to underlying cellular mechanisms. It is common to handle this variability by simply averaging over the data and finding a single set of model parameters that best fits the averaged data. The “populations of models” approach allows deterministic models to reflect the inherent variability in biological data through identification of not just the single best parameter set but a population of parameter sets such that the output of the group of models displays the same heterogeneity as the population being modeled [9, 11–16]. The problem of constructing populations of deterministic models and identifying distributions of model input parameters from stochastic observations from multiple individuals in a population is known as the stochastic inverse problem (SIP). State-of-the-art methods for solving SIPs apply Bayesian inference techniques, including Markov chain Monte Carlo (MCMC) sampling, and are limited to finding a distribution for a single set of observations [11, 17–19]. To draw inferences about a new target dataset, the SIP would have to be solved again. We have recently proposed an alternative approach to solving SIPs, using generative adversarial networks (GANs), that enables *amortized inference*— i.e, the trained GAN can be reused on many target datasets without retraining [20]. GANs are a deep learning paradigm involving two artificial neural networks that compete with each other in a minimax game. The *generator* network attempts to produce fake samples that are as similar as possible to a distribution of real samples, and the *discriminator* network tries to distinguish fake samples from real samples. Since being introduced in 2014, GANs have garnered significant interest across a wide range of fields, including applications in image processing, cybersecurity, cryptography, and neuroscience [21–23]. Several extensions of GANs have been developed to address particular tasks [24–26]. To solve SIPs, we use a conditional GAN (cGAN) structure [27] where the generator is trained with parameter sets *X* conditioned on the output features *Y* of a mechanistic model.

In this paper, we wish to solve an SIP to identify which ion channels are responsible for the altered excitability properties of hippocampal neurons in mouse models of amyloidopathy and tauopathy. The data for the SIP of interest in this paper are voltage traces recorded from hippocampal CA1 neurons in 12-month-old rTg4510 mice expressing pathogenic tau (Tamagnini et al., unpublished data), 24-month-old PDAPP mice overexpressing amyloid beta [3], and age-matched wildtype littermate controls for each transgenic phenotype. From these traces, we extract several electrophysiological features (such as action potential peak, width, and threshold) that capture the excitability properties of the cells. Neuronal excitability can be simulated using the conductance-based modeling formalism originally developed by Hodgkin and Huxley [28]. The mechanistic model for the SIP of interest in this paper is a conductance-based model of CA1 neurons that has been shown to be capable of reproducing key electrophysiological features of recorded voltage traces [6, 29]. By solving this SIP, we will map the features of the recorded voltage traces in the AD mutant and wildtype mice to the parameter space of the CA1 model. Our goal is to use the resulting parameter distributions to infer which ion channel conductances are disrupted in the amyloidopathy and tauopathy mice compared to their age-matched wildtype controls, and which conductances change with age in the wildtype mice.

The remainder of the paper is organized as follows. In Section 2, we describe the experimental data and the features we extract from the recorded action potentials and hyperpolarization traces. In Section 3, we introduce a biophysical model of CA1 pyramidal neurons and our initial optimizations of the model parameters using differential evolution. In Section 4, we give a brief description of GANs and cGANs and illustrate our parameter inference methodology using the Rosenbrock function as a toy model. In Section 5, we train the cGAN on output of the CA1 model and then present it with synthetic target data. We show that cGAN outperforms a benchmark MCMC method on a relatively simple parameter inference task. We then validate its ability to accurately infer complex parameter distributions through a series of tests with synthetic target data. In Section 6, we present the trained cGAN with real target data and use the inferred parameter distributions to identify which ionic conductances are affected by age, amyloidopathy, and tauopathy. We conclude with a discussion of alternative methods and future work in Section 7.

## 2 Experimental Data and Feature Extraction

The experimental data we use consists of patch-clamp recordings made from hippocampal CA1 neurons associated with two previous studies involving mouse models of Alzheimer’s disease. In Tamagnini et al. [3], CA1 current-clamp recordings were obtained from transgenic PDAPP mice exhibiting amyloidopathy. In unpublished data, Tamagnini et al. obtained CA1 current-clamp recordings from transgenic rTg4510 mice exhibiting tauopathy. In this paper, we use voltage traces from *n* = 30 cells of 24-month-old PDAPP mice (and *n* = 19 cells from their age-matched WT littermate controls) and *n* = 26 cells of 12-month-old rTg4510 mice (and *n* = 26 cells from their age-matched WT littermate controls).

To characterize the excitability of these cells, we focused on the properties of the first action potential (AP) elicited in response to a square depolarizing current pulse (300 pA, 500 ms; Fig. 1A) and on the electrotonic properties of the plasma membrane measured upon the membrane potential exponential decay in response to a square hyperpolarizing current pulse (−100 pA, 500 ms; Fig. 1B). To account for the biasing effect of cell-to-cell variability of the membrane potential over the excitability properties, all recordings were made from a starting membrane potential of *V*_*m*_ = −80 mV. This *V*_*m*_ value was obtained via the constant injection of a biasing current. To summarize the behavior of these voltage traces, we defined 9 features associated with the APs and 4 features associated with the membrane hyperpolarization.

**Fig. 1.**
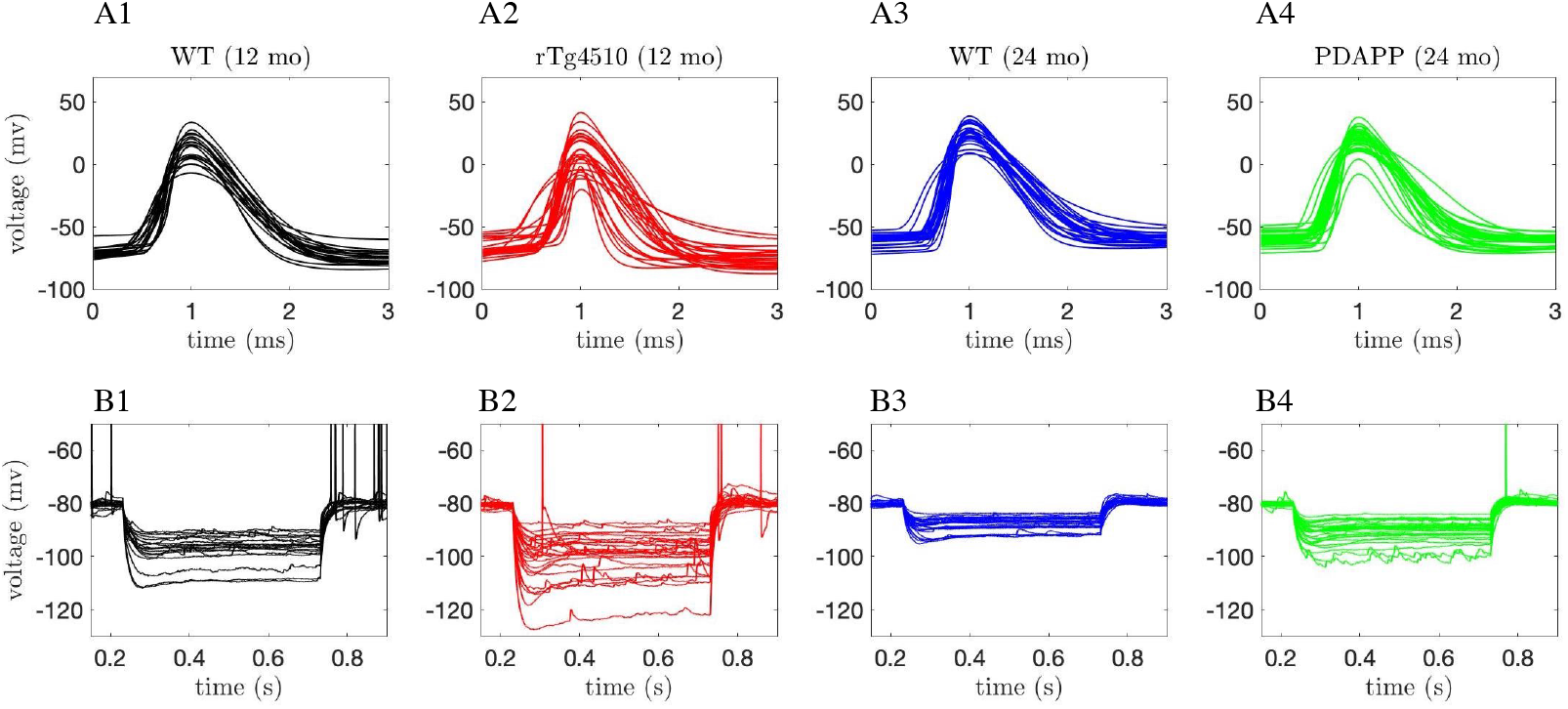
Experimental recordings. Waveforms of the first action potential in response to depolarizing current pulses (**A1-A4**) and voltage traces in response to hyperpolarizing current pulses (**B1-B4**) injected into CA1 pyramidal neurons for 4 different categories of mice: wildtype (WT) 12-month-old mice (black traces, **A1-B1**), tau mutant (rTg4510) 12-month old mice (red traces, **A2-B2**), WT 24-month old mice (blue traces, **A3-B3**), and amyloid beta mutant (PDAPP) 24-month-old mice (green traces, **A4-B4**).

The AP features are illustrated in Figs. 2A-B and are defined as follows:

**Fig. 2.**
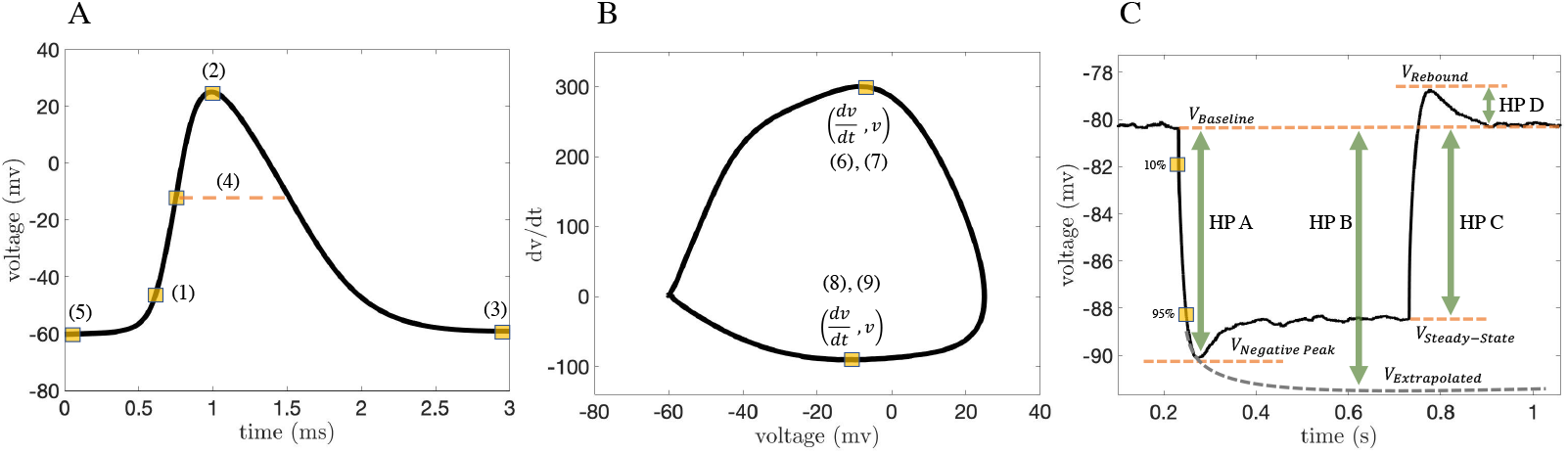
Schematic of feature extraction. **(A-B)** Action potential features: (1) AP threshold, (2) AP peak, (3) AP trough, (4) AP width, (5) AP min voltage before the pulse, (6) AP max positive rate of rise, (7) AP voltage at max positive rate of rise, (8) AP max negative rate of rise, (9) AP voltage at max negative rate of rise. **(C)** Hyperpolarization features: (10) HP A - voltage at negative peak and baseline differences, (11) HP B - voltage at exponential fit and baseline differences, (12) HP C - voltage at steady state and baseline differences and (13) HP D - voltage at rebound and baseline differences.

1. *AP threshold:* voltage at 10 percent of the AP max positive rate of rise (feature 6)
2. *AP peak:* maximum value of the voltage trace
3. *AP trough:* minimum value of the voltage in the 2 ms time interval after the AP peak
4. *AP width:* duration of time that the voltage is above the AP voltage at max positive rate of rise (feature 7)
5. *AP min voltage before the pulse:* minimum voltage in the 1 ms interval before the AP peak
6. *AP max positive rate of rise:* maximum value of *dV/dt* in the 3 ms time interval around the AP peak (i.e. 1 ms before and 2 ms after the peak)
7. *AP voltage at max positive rate of rise:* voltage value at the AP max positive rate of rise
8. *AP max negative rate of rise:* minimum value of *dV/dt* in the 3 ms time interval around the AP peak (i.e. 1 ms before and 2 ms after the peak)
9. *AP voltage at max negative rate of rise:* voltage value at the AP max negative rate of rise. The membrane hyperpolarization features are illustrated in Fig. 2C and are defined as follows:
10. *HP A* - voltage at negative peak and baseline differences
11. *HP B* - voltage at exponential fit and baseline differences
12. *HP C* - voltage at steady state and baseline differences
13. *HP D* - voltage at rebound and baseline differences.

We note that these features were chosen to try to capture as much of the behavior of the voltage traces in as few features as possible. Increasing the dimensionality of the feature space can reduce the accuracy of cGAN training if the additional features are not sufficiently informative.

We then calculated these features for the voltage traces from PDAPP, rTg4510, and WT mice (see Fig. 3 for the AP features, and Fig. 4 for the hyperpolarization features). Despite the large amount of variability within each category, for some features clear differences are observed across categories. For example, AP peak appears to be reduced in PDAPP mice compared to their WT controls (Fig. 3 top middle panel) and AP width appears to be reduced in rTg4510 mice compared to their WT controls (Fig. 3 middle left panel).

**Fig. 3.**
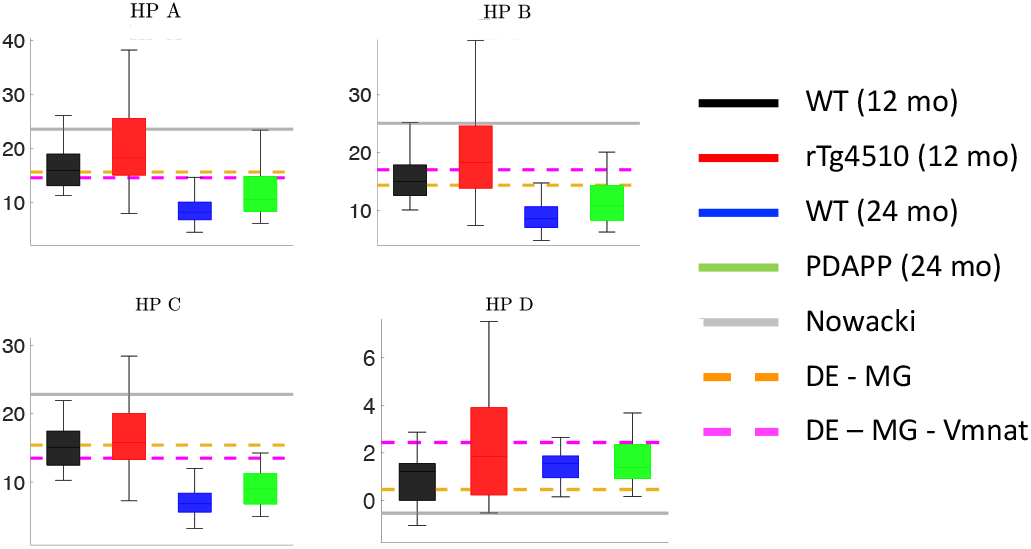
Action potential features in experimental data and initial models. Box and whisker plots of the action potential (AP) features extracted from the experimental data and the biophysical CA1 model with three different parameter sets: (1) the parameters in the original Nowacki et al. paper ([29], solid gray lines), (2) parameters obtained using differential evolution, optimizing only the maximal conductance parameters (DE-MG, dashed orange lines), and (3) parameters obtained using differential evolution, optimizing the maximal conductances and the half-activation voltage of the transient sodium current (DE-MG-Vmnat, dashed magenta lines).

**Fig. 4.**
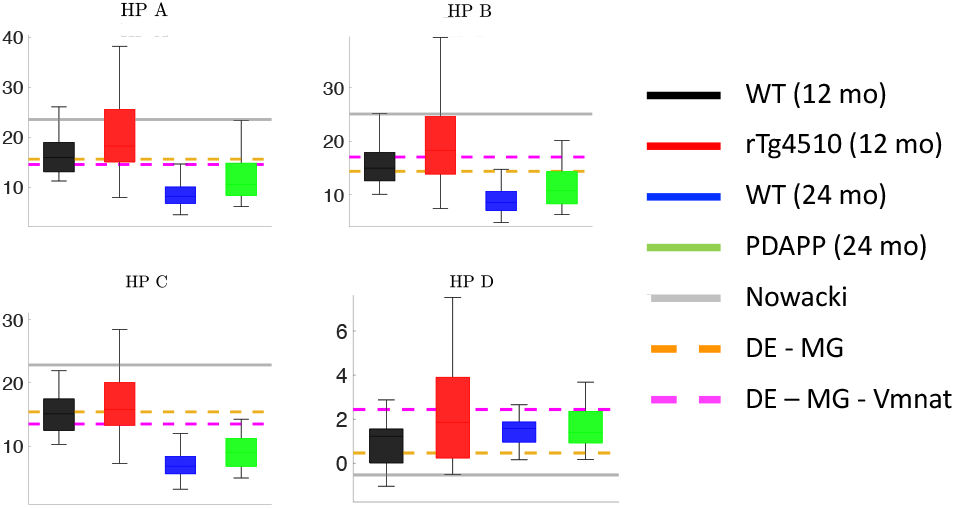
Membrane hyperpolarization features in experimental data and initial models. Box and whisker plots of the membrane hyperpolarization features extracted from the experimental data and the biophysical CA1 model with three different parameter sets: (1) the parameters in the original Nowacki et al. paper ([29], solid gray lines), (2) parameters obtained using differential evolution, optimizing only the maximal conductance parameters (DE-MG, dashed orange lines), and (3) parameters obtained using differential evolution, optimizing the maximal conductances and the half-activation voltage of the transient sodium current (DE-MG-Vmnat, dashed magenta lines).

## 3 Biophysical Model

### CA1 pyramidal neuron model

Conductance-based modeling to describe the electrical activity of neurons was introduced by Hodgkin and Huxley in 1952 to explain the ionic mechanisms underlying the initiation and propagation of action potentials (APs) in the squid giant axon [30]. Nowacki et al [29] developed a conductance-based model of CA1 pyramidal neurons that includes the following ionic currents: two *Na*^+^-currents, one transient 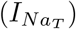 and one persistent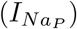; two *Ca*^2+^-currents, one T-type 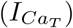 and one high-voltage activated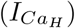; and three *K*^+^-currents, delayed rectifier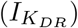, M-type 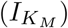, and leak (*I*_*L*_). The dynamics of the membrane potential *V* and ionic gating variables *x* are governed by the following system of ordinary differential equations:

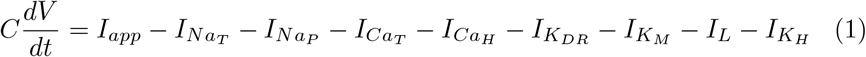

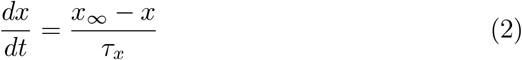

where:

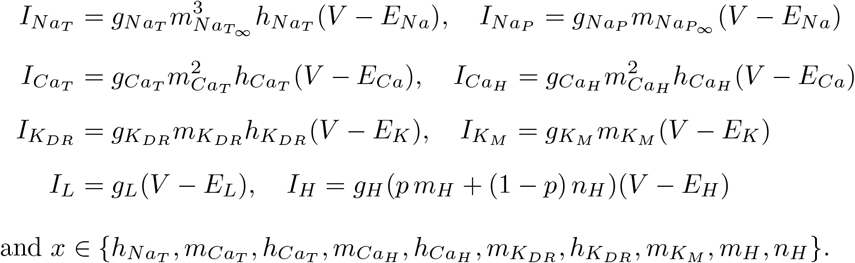

and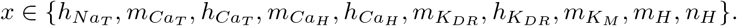

The ionic currents *I* are described by Ohm’s Law with maximal conductance parameters *g* and reversal potentials *E*. The steady-state activation and inactivation functions *x*_∞_ for all gating variables, including *m*_*NaT*_ and *m*_*NaP*_, are given in Boltzmann form:

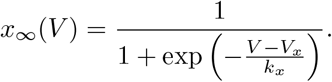

The time constants *τ*_*x*_ for all gating variables are fixed parameters, except for *h*_*NaT*_, for which the time constant is a voltage-dependent function:

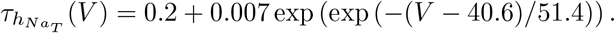

First, we simulated the pyramidal neuron model with the parameter values provided in Nowacki et al [29] (Supplementary Table 1) and calculated feature values based on the model output (i.e. the simulated voltage trace). For certain features, the model’s feature value is outside the range of the feature values observed in the experimental data (solid gray lines in Figs. 3 and 4). For example, the AP threshold and AP peak in the model are significantly more depolarized than the AP thresholds and peaks seen in these CA1 neurons (Fig. 3 top left and top middle panels).

Thus, we used stochastic global optimization to find model parameters that produce model output with feature values consistent with the experimentally observed feature values. Specifically, we used differential evolution (DE), a population-based search technique first introduced by Storn and Price [31, 32].

The objective function for the DE algorithm was to minimize the sum of squares error between the simulated voltage trace and the average voltage trace for each category (PDAPP, rTg4510, WT 12 and 24 month) across all four categories. More details on our implementation of the DE algorithm are provided in the Supplementary Methods.

Initially, we chose to hold all the reversal potentials and gating variable parameters at their original Nowacki et al. values, so the only free parameters for DE to optimize were the 8 maximal conductances. The model with optimized maximal conductances produced model output with feature values more consistent with the range of the feature values in the experimental data (dashed orange lines in Figs. 3 and 4). However, this model’s AP threshold was still significantly more depolarized than in the data (Fig. 3 top left panel). We used a variance-based sensitivity analysis (Sobol’s method) to determine which model parameters affect the model’s AP threshold, and found that the half-activation of the transient sodium current (*V*_*mNaT*_) has the largest effect (see Supplementary Methods for a description of our sensitivity analysis procedure). We then ran DE again, this time with *V*_*mNaT*_ as a free parameter in addition to the maximal conductances. The model with optimized *V*_*mNaT*_ produced model output with feature values within the range of the feature values in the experimental data, including the AP threshold (dashed magenta lines in Figs. 3 and 4). Furthermore, the action potential and membrane hyperpolarization voltage traces produced by this model agree well with the experimental voltage traces themselves, as the model output appears to lie in the middle of the four voltage traces obtained by averaging the traces within each of the four categories (Fig. 5).

**Fig. 5.**
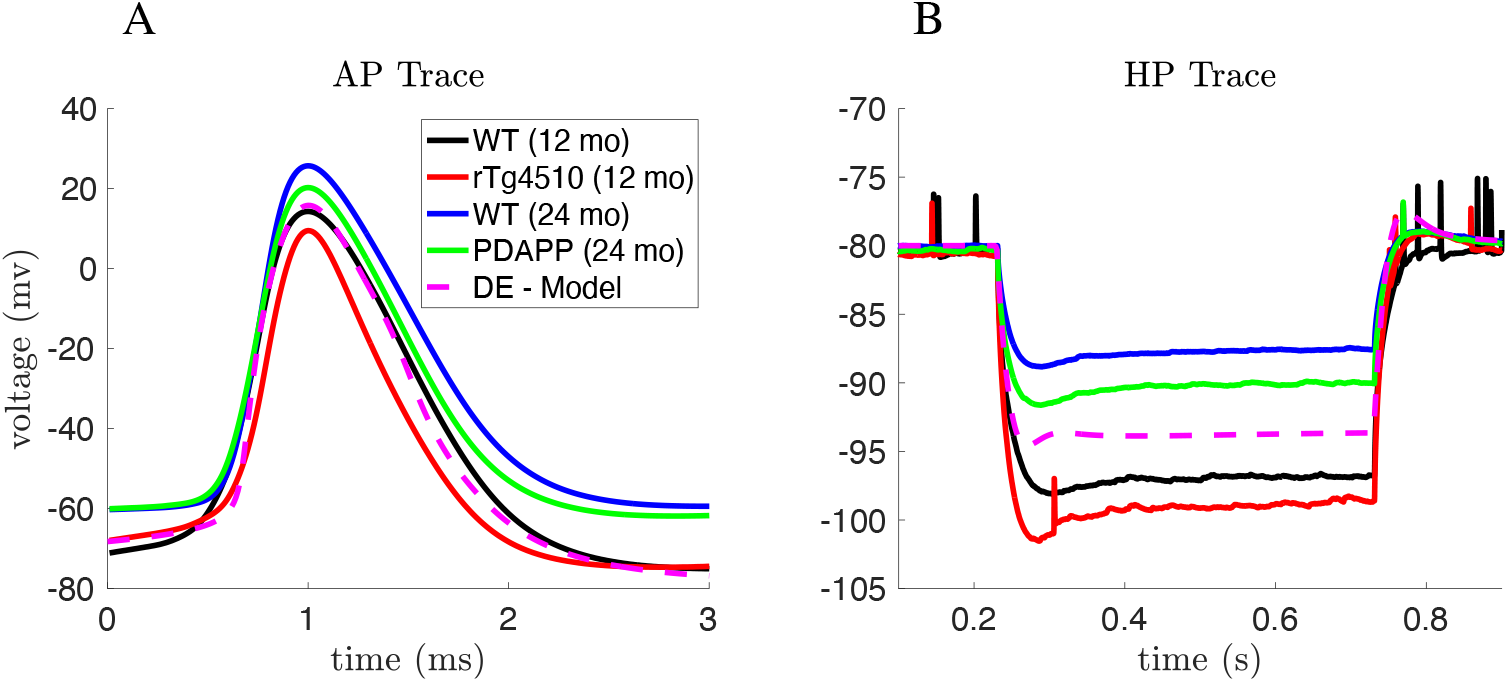
Average AP and hyperpolarization voltage traces from experimental data and optimized model. The DE-Model shown here is the same model that was labeled DE-MG-Vmnat in Fig. 4, and was obtained by minimizing the least square error between the DE-Model output and the average AP and hyperpolarization traces across all 4 categories. **(A)** Mean of the AP waveforms in the experimental data for each category, and the AP waveform simulated using the optimized DE model (magenta). **(B)** Mean of the membrane hyperpolarization traces in the experimental data for each category, and the hyperpolarization trace simulated using the optimized DE model (magenta).

We refer to the parameter set obtained through DE with *V*_*mNaT*_ and maximal conductances as free parameters as the “default” model parameters. We will use these parameters to set parameter bounds when creating the training dataset for cGAN. The default parameters are provided in Supplementary Table 1.

## 4 Parameter Inference Methodology: conditional Generative Adversarial Networks

### 4.1 Generative Adversarial Networks

Generative Adversarial Networks (GANs) are an example of generative models in machine learning. Since GANs have a deep neural network architecture we can classify them as deep learning models. The application we are interested in here is to build an inverse surrogate model for mapping the output of a mechanistic model into its corresponding region in parameter space. More precisely, the goal is to map the density of observed data (𝒫_*Y*_) to a *coherent* density *α*_*X*_ of the model input parameter space. A distribution *α*_*X*_ is coherent if: (1) upon sampling from *α*_*X*_ and applying the mechanistic model, the estimated density in output space 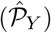 satisfies 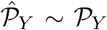, and (2) *α*_*X*_ covers all possible solutions in the range described by the prior in input space (𝒫_*X*_). In this section, we will first introduce standard GANs, and then move on to conditional GANs (cGANs) that are designed to incorporate conditional distributions into GANs.

The basic GAN consists of two artificial neural networks, a generator *G* and a discriminator *D*, that compete with each other (Fig. 6). The Generator tries to produce fake samples that are as close as possible to real samples that come from some distribution, and the Discriminator tries to distinguish real samples from fake samples. Training the GAN is an iterative process through which *G* gets better at fooling *D*, and *D* gets better at identifying fake samples. The first step in training the GAN is to create a training dataset (referred to as real samples) by simulating the mechanistic model with parameter sets *x* drawn from a uniform distribution *P*_*data*_(*x*). Then we initialize the Generator with a base distribution *P*_*z*_(*z*) which is a random variable with typically a Gaussian distribution. These parameter sets *G*(*z*) ∼ *P*_*G*_, referred to as fake samples, are passed to the Discriminator along with the real samples. For a given sample *x* or *G*(*z*), the Discriminator outputs a probability *ŷ* = *D*(*x*) or *ŷ* = *D*(*G*(*z*)), referred to as a reconstructed label, indicating whether it thinks the sample is real (*ŷ >* 0.5) or fake (*ŷ<* 0.5). If *D* is correct (i.e. *ŷ* = *D*(*x*) *>* 0.5 or *ŷ* = *D*(*G*(*z*)) *<* 0.5), then the weights and biases (*ω, β*) of the *D* network will remain fixed, but (*ω, β*) of the *G* network will be adjusted through backpropagation (referred to as fine-tuning in Fig. 6). If *D* is incorrect (i.e. *ŷ* = *D*(*x*) *<* 0.5 or *ŷ* = *D*(*G*(*z*)) *>* 0.5), then (*ω, β*) of *D* are adjusted while (*ω, β*) of *G* remain fixed. In practice, convergence of the generator and discriminator one at a time would not only be time consuming, but also lead to instability due to a vanishing gradient for the loss function of the generator. Therefore, in this case the weights are adjusted after computing the loss function from the outputs of *D* and *G* over each mini-batch. As a result, both the generator and the discriminator are being trained simultaneously, and they are converging gradually.

**Fig. 6.**
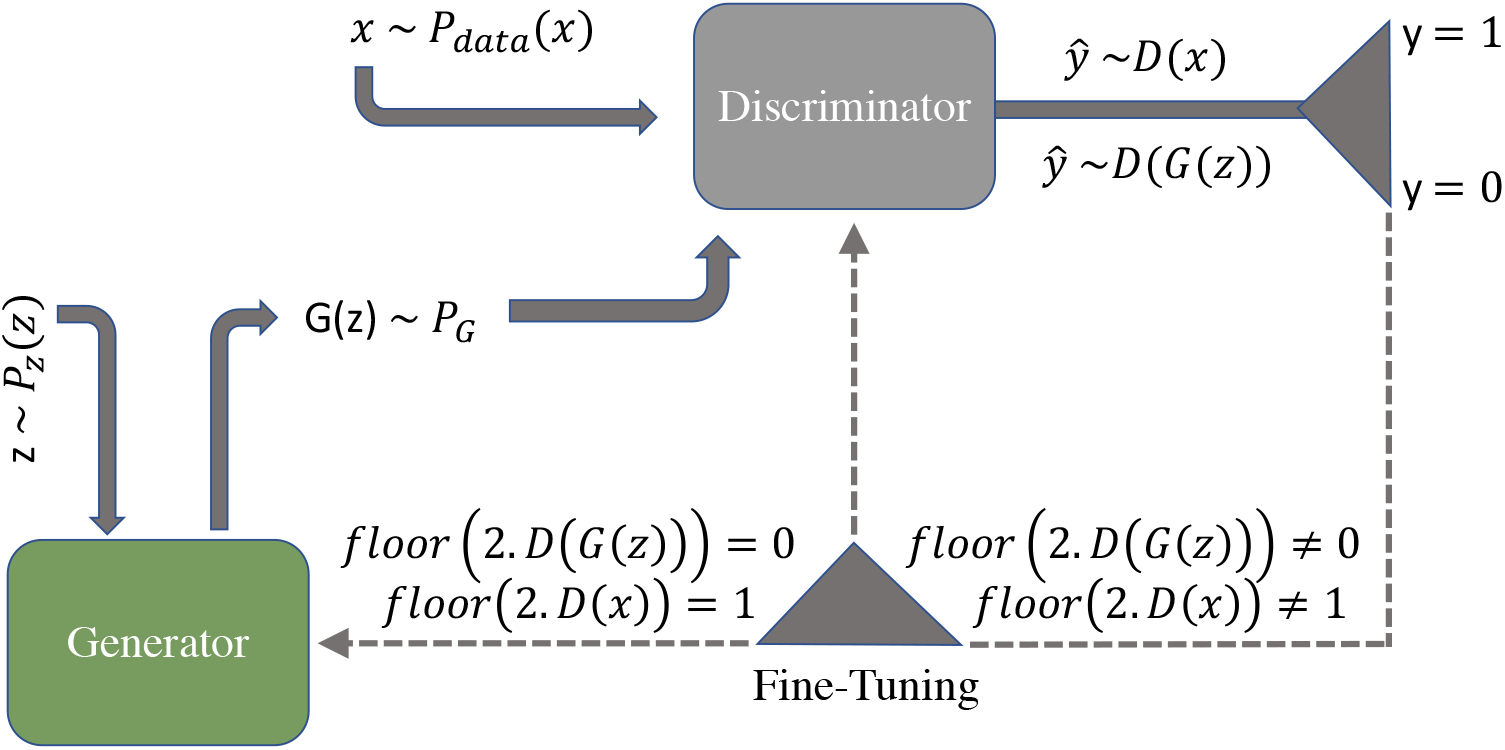
Schematic of a generative adversarial network (GAN). The Generator (*G*) and Discriminator (*D*) are two neural networks that compete with each other during the training process, with the end result being that *G* can produce fake samples from a distribution that matches the distribution of the real samples. *D*(*x*) and *D*(*G*(*z*)) provide the reconstructed label *ŷ* (i.e. the output of the Discriminator network), which represents the probability of the sample being real rather than fake. If *ŷ* is greater than (less than) 0.5, the Discriminator classifies the sample as real (fake). If *D* is correct (incorrect), then the weights of *G* (*D*) are fine-tuned through backpropagation.

#### Derivation of the objective function for the GAN

*Cross-entropy* is a measure from the field of information theory which calculates the difference between two probability distributions and is defined by the following equation:

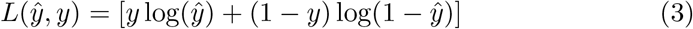

Where *ŷ* is the reconstructed label *D*(*x*) or *D*(*G*(*z*)) and *y* is the actual label for the sample. Therefore, the corresponding label for a real sample (i.e. a sample coming from *P*_*data*_(*x*)) is *y* = 1 and the reconstructed label (i.e. the output of the Discriminator) is *ŷ* = *D*(*x*). By substituting these labels for the real samples into the equation (3) we can get:

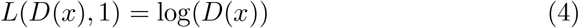

Likewise the data coming from the Generator has the real label *y* = 0 and the reconstructed label is *ŷ* = *D*(*G*(*z*)), therefore, by substituting these expressions for the fake samples into Eqn. (3) we end up with:

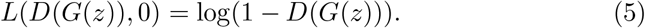

Panels A and B in Figure (7) are the visualization of Eqns. (4) and (5), respectively. Since the output of the Discriminator is a probability, in both of these panels we only consider the region between zero and one on the *x*−axis (which represents *D*(*x*) or *D*(*G*(*z*))). In panel A, log(*D*(*x*)) is an increasing function of *D*(*x*). If we have a strong Discriminator, then we expect *D*(*x*) ∼ 1 for a real sample. In panel B, the *x*−axis represents *D*(*G*(*z*)), and the expectation for a strong Discriminator would be *D*(*G*(*z*)) ∼ 0 for a real sample. As these two points are close to the maximum of both Eqns. (4) and (5), the objective function for the Discriminator is:

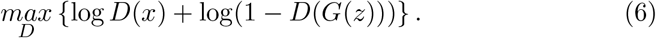

The same logic also holds in the case of having a strong Generator, the only difference here is that we are going to minimize Eqn. (5), which corresponds to the right panel of Fig. 7. Having a very strong Generator means it is able to fool the Discriminator easily. This means the Discriminator will erroneously return a high probability even for a fake sample, i.e. *D*(*G*(*z*)) ∼ 1. This point is the minimum value of Eqn. (5). Thus, the objective function for the Generator is:

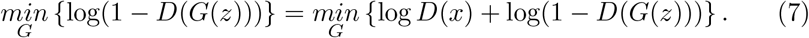

In order to obtain a single objective function for the GAN, we combine Eqns. (6) and (7) and take the expectation over the whole dataset. The resulting objective function for the GAN, which is inspired by the cross-entropy loss, is given by:

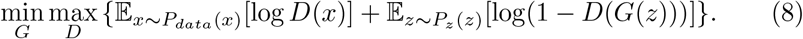

In practice, at the beginning of the training process the Generator is not strong enough and the output of the Generator is very different from the training dataset. Thus, the Discriminator can easily distinguish the real and the fake samples. This causes log(1 − *D*(*G*(*z*))) to saturate (see Fig. 7B), and the gradient does not provide any information as it is almost zero. Therefore, as is mentioned in [21], instead of minimizing log(1−*D*(*G*(*z*))) we can minimize the log(1 − *D*(*G*(*z*))) − log(*D*(*G*(*z*))). This will help ensure we have a more stable loss function during the training process.

**Fig. 7.**
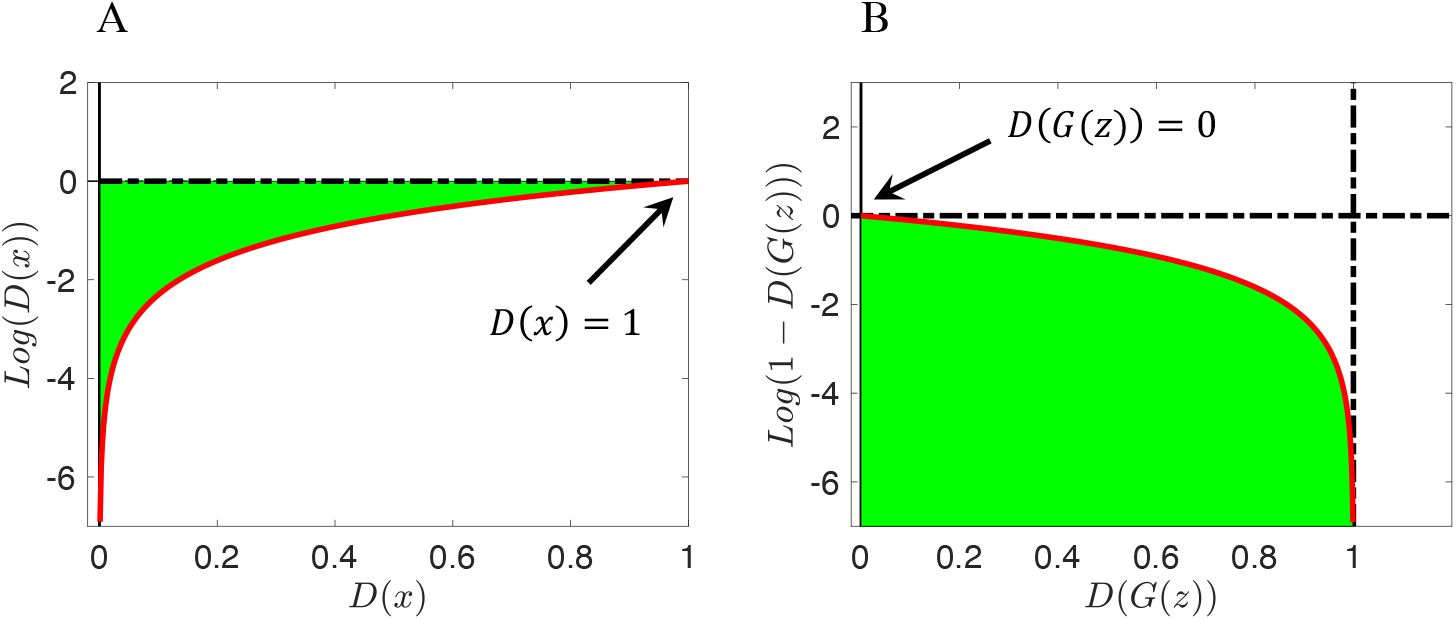
Visualization of the components of the GAN objective function (Eqn. (8)). **(A)** Discriminator loss function (Eqn. (4)). *D*(*x*) is the probability that *x* came from the real data rather than *P*_*g*_. **(B)** Generator loss function given (Eqn. (5)). *D*(*G*(*z*))) is the probability that *G*(*z*) came from *P*_*g*_ rather than *P*_*data*_. The green shading identifies the possible regions for these two probabilities.

### 4.2 Conditional Generative Adversarial Networks

A standard GAN could be trained to produce samples of parameter sets, samples of feature sets, or even samples of combined parameter-feature sets. However, it is not able to produce samples of parameter sets corresponding to a set of given feature values. To accomplish this, we employ conditional GANs (cGANs) [27], where features extracted from the output of the mechanistic model are passed as a condition to the Generator. The parameter samples produced by the Generator, augmented with the features it was provided as a condition, are then passed to the Discriminator. Ground truth parameters, with their corresponding features, as a condition, are also passed to the Discriminator. During the training process, the Generator learns how to produce samples in parameter space that are similar to the ground truth parameters for a given set of features.

The overall structure of the cGAN is similar to the basic GAN model. However, the main difference is that the input into both the Generator (*G*) and Discriminator (*D*) are augmented by the conditional variable *Y* as described in Fig. 8. The objective function of the cGAN is:

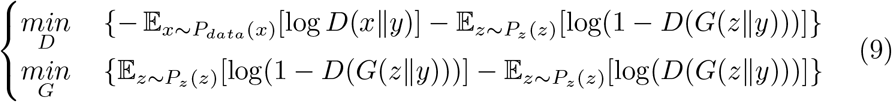

**Fig. 8.**
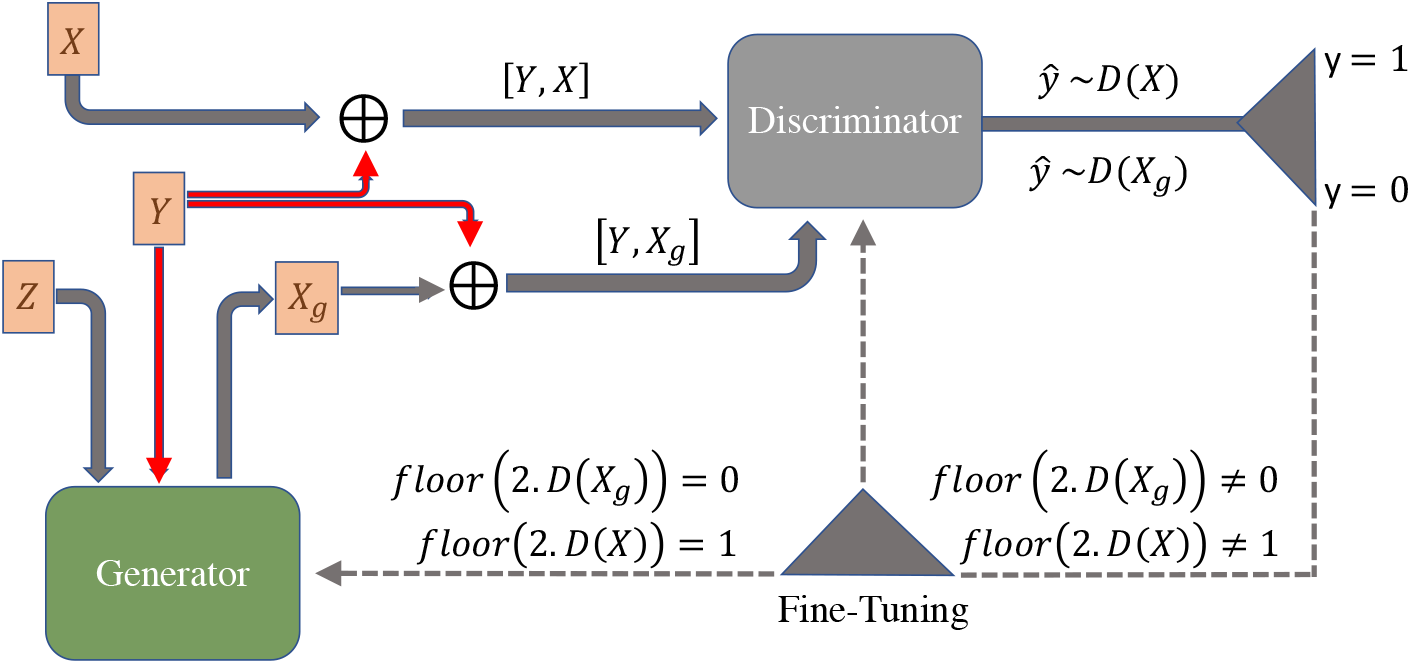
Schematic of a conditional GAN (cGAN). Features of the training dataset (*Y*) are passed as a condition into the Generator (*G*) - already initialized with a Gaussian distribution (*Z*) - in order to produce some samples *X*_*g*_ given that condition [*Y, X*_*g*_]. This output, along with real samples *X* augmented with their output features [*Y, X*], are passed into the Discriminator (*D*). If the Discriminator’s output is close to zero (one), it means the Discriminator has assigned a low (high) probability of that sample being real.

### 4.3 Illustration of cGAN training process

We used the Rosenbrock function (Eqn. 10) as a toy mechanistic model to illustrate the training process for cGAN:

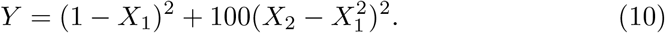

In this example, we have only two input parameters (*X*_1_ and *X*_2_) and only one feature (*Y*, the output of the Rosenbrock function). Thus, with a fixed output as a condition, cGAN converges to the ground truth region in the parameter space after a few training epochs. Figure (9) provides a visual summary of the training. We used Jensen Shannon Divergence (JSD), a measure of the similarity between two probability distributions, as our stopping criterion (see Supplementary Methods for more details on our JSD computation). When the JSD measure for a fixed epoch number approaches zero, then we stop training after that epoch. If one continues training after this stopping point, the training process destabilizes and the JSD measure starts increasing (Fig. 9 bottom right panel). This phenomenon occurs due to the vanishing gradient problem.

**Fig. 9.**
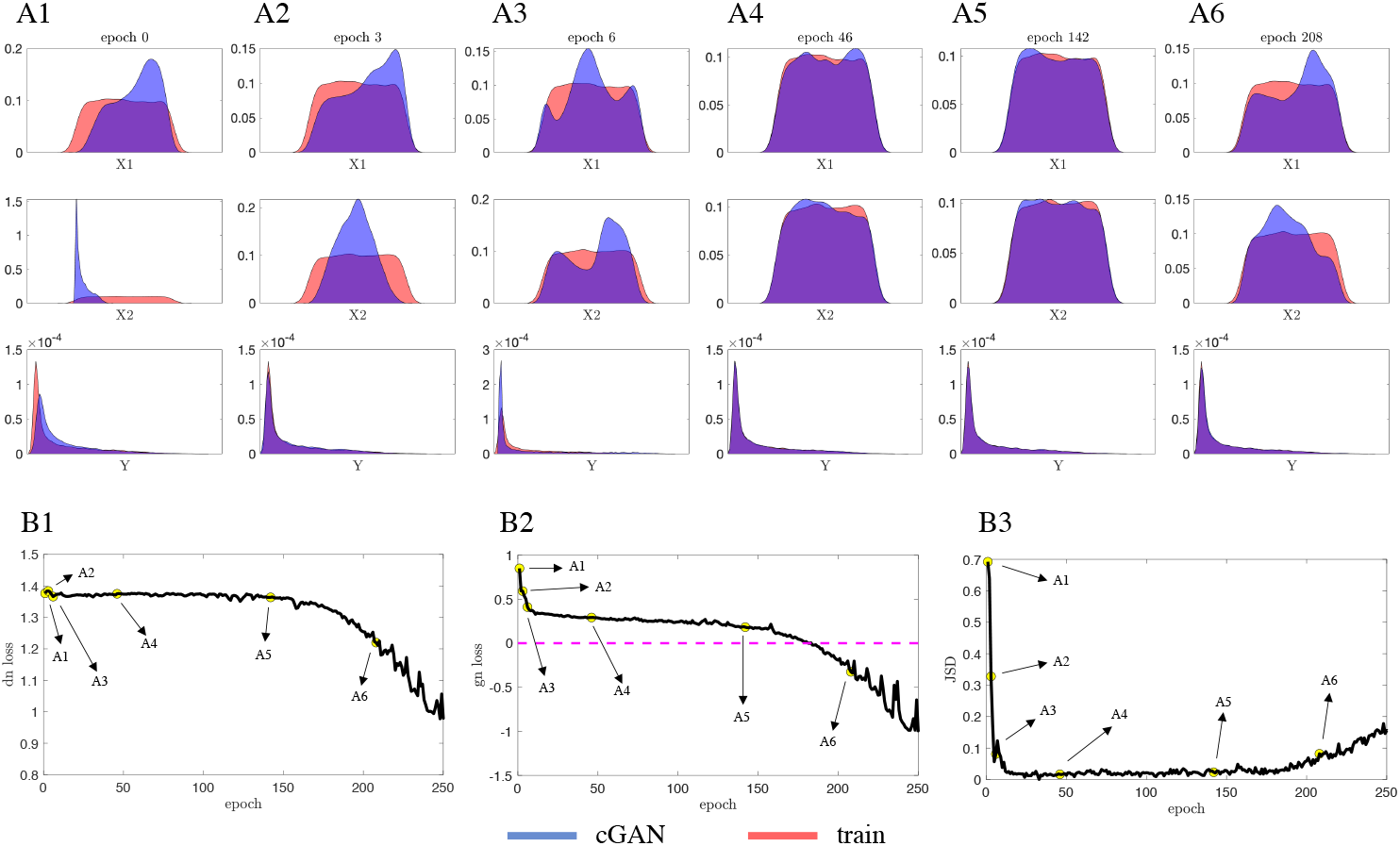
Illustration of the cGAN training process on the Rosenbrock function (Eqn. 10). Over the course of the training process, both the discriminator and the generator improve. The discriminator gets better at distinguishing between real (coming from *P*_*data*_) and fake (coming from *P*_*g*_) samples, and for a fixed discriminator, the generator gets better at fooling the discriminator. **(A1-A6)** KDE plots of the cGAN samples (blue) and the training data (red) for parameters *X*1, *X*2, and feature *Y* at epochs 0, 3, 6, 46, 142, and 208. At epoch 142 (A5), the cGAN distributions are good approximations of the training distributions, but by epoch 208 (A6) the cGAN distributions are no longer good approximations. **(B1-B3)** The discriminator loss (B1), the generator loss (B2), and the JSD measure between the ground truth parameters versus estimated parameters (B3) as a function of epoch number. The point labeled A5 in panel B3 represents the JSD stopping criterion: once the JSD starts to increase we stop training the cGAN.

## 5 cGAN Training on Biophysical Model and Validation on Synthetic Target Data

Recall that our main goal is to use cGAN to map voltage traces recorded from WT and Alzheimer’s mutant mice to the parameter space of our CA1 model. To enable cGAN to learn the mapping from electrophysiological features to the CA1 model parameter space, we will create a training dataset consisting of features calculated from CA1 model simulations with randomly chosen parameter values. Since we are primarily interested in how the maximal conductances of the various ion channels present in CA1 pyramidal neurons are affected by aging and amyloidopathy/tauopathy, we will only vary the maximal conductance parameters in our training dataset and keep the gating variable kinetic parameters and the reversal potentials fixed at their default values. However, it may be that some of the maximal conductances do not have a large effect on the output features of interest. To explore this possibility, before creating a training dataset, we first conducted Sobol sensitivity analysis to see how each of the 8 maximal conductance parameters affect the features. We found that 3 of these conductances, *g*_*NaP*_, *g*_*CaT*_, and *g*_*L*_ have a small effect on the features compared to the other 5 conductances (Fig. 10). From a biological standpoint, these 3 conductances are unlikely to play a major role in determining the AP features for the following reasons: (1) persistent sodium current (*I*_*NaP*_) is likely to be much smaller than transient sodium current (*I*_*NaT*_), (2) T-type calcium (*I*_*CaT*_) is likely to be much slower than *I*_*NaT*_, and (3) the leak current (*I*_*L*_) primarily affects resting membrane potential rather than AP dynamics. Therefore, when we create the training dataset, we keep those 3 conductances fixed at their default values, and only vary 5 conductances: *g*_*NaT*_, *g*_*CaH*_, *g*_*KDR*_, *g*_*KM*_, and *g*_*H*_ .

**Fig. 10.**
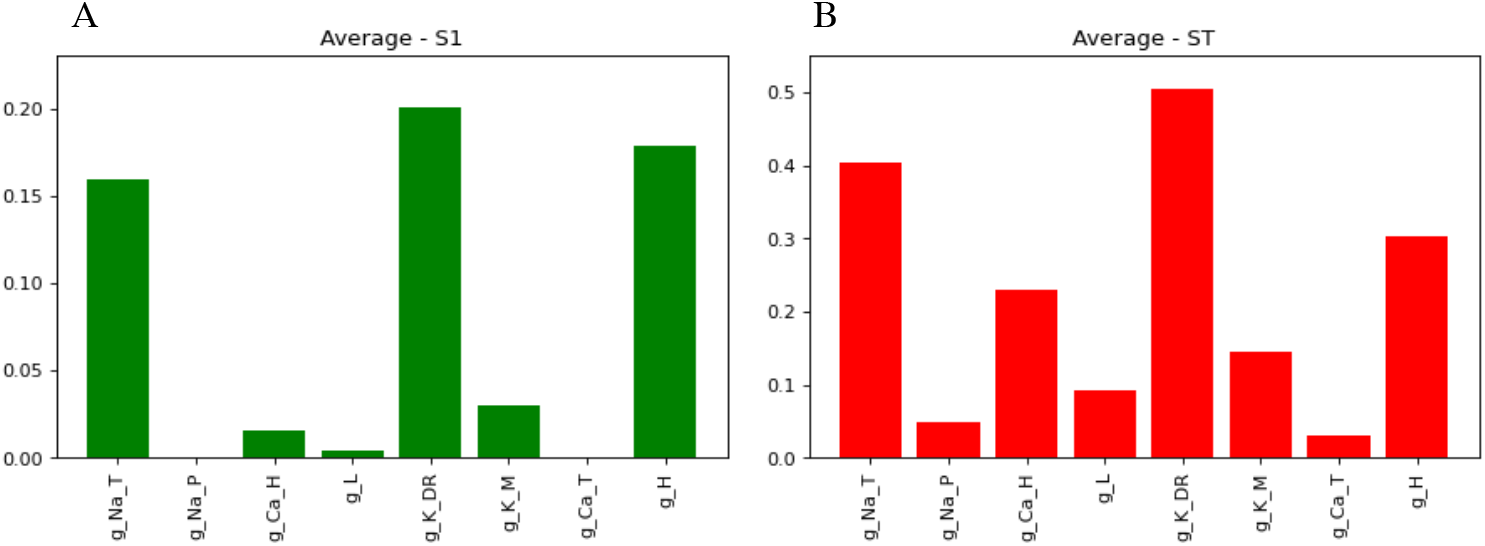
Dimensionality reduction using variance-based (Sobol) sensitivity analysis. The height of the bars represent how sensitive the AP and hyperpolarization features are to each parameter. **(A)** S1: average first-order index across feature space. **(B)** ST: average total-effect index across feature space.

For the training dataset, we performed 3 million simulations of the CA1 model with these 5 conductances drawn from uniform distributions with upper and lower bounds at *±*100% of their default values. We then calculated the feature values for these simulations, and trained the cGAN with the parameters *X* conditioned on the features *Y*. Once the cGAN was trained, we passed the features for a subset of the training dataset (10,000 simulations) to the trained cGAN and asked it to produce samples (i.e. parameter sets) for those features. We then simulated the CA1 model with the parameter sets from the cGAN and calculated the features from these simulations. To compare the distributions of features from the training dataset and from the cGAN samples, we plotted Kernel Density Estimates (KDEs) for each feature and scatter plots for each pair of features (Fig. 11A). These plots show that the cGAN samples produced features that were very consistent with the features in the training data. We also plotted KDEs and pairwise contour plots for each parameter, which show that the distributions of parameters produced by the cGAN are similar to the parameter distributions in the training dataset (Fig. 11B). This visual conclusion was confirmed by calculating the Jensen Shannon Divergence. The JSD between the cGAN samples and training data on the joint distribution of parameters and features was approximately zero (1.11*×*10^−14^).

**Fig. 11.**
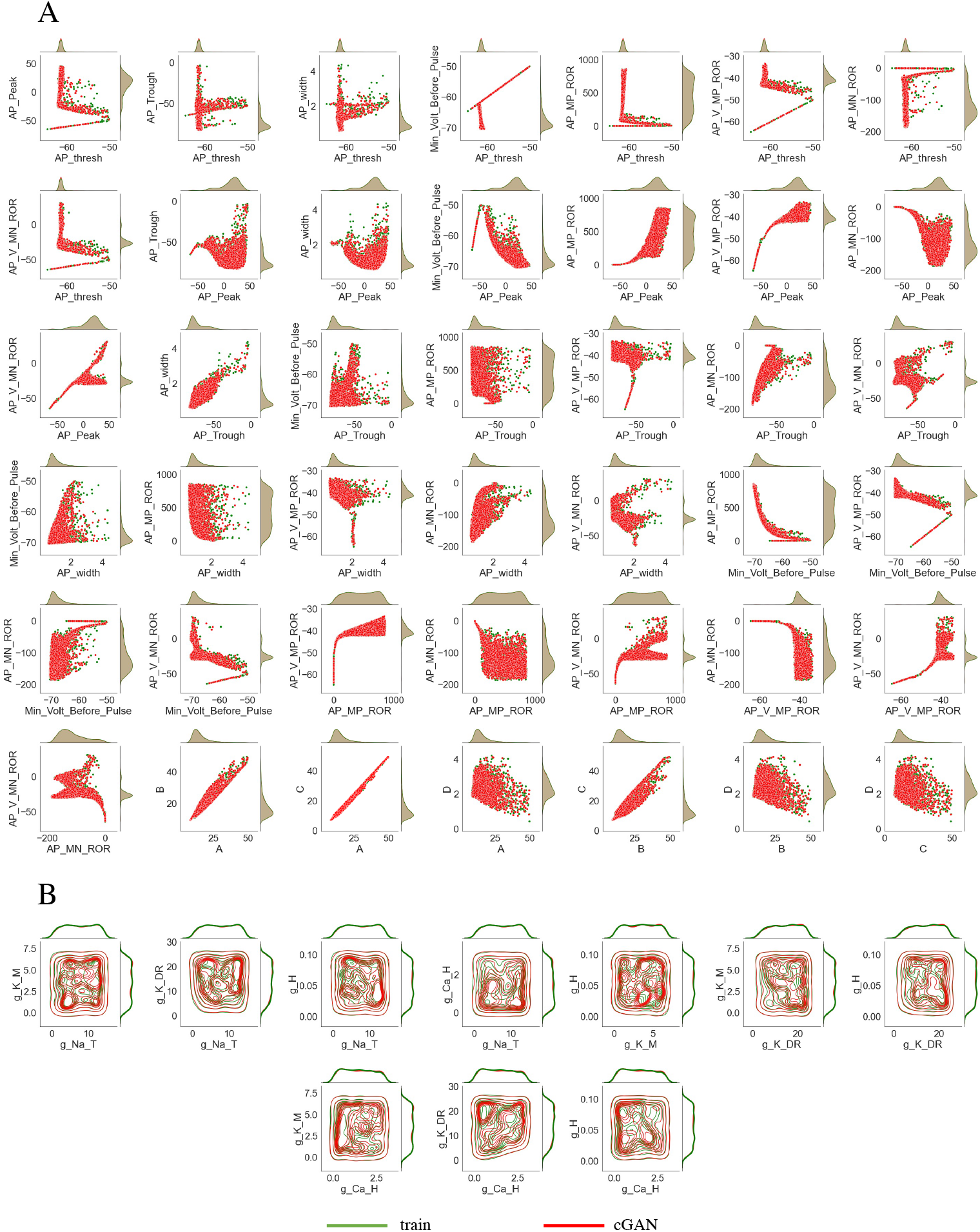
Comparison of cGAN samples and training dataset. **(A)** Feature space: scatterplots (center of panel) and KDE plots (top and right of each panel) with cGAN samples in red and the training dataset in green. **(B)** Parameter space: contour plots (center of panel) and KDE plots (top and right of each panel) with cGAN samples in red and the training dataset in green. In both (A) and (B), the KDE plots for the cGAN samples are nearly identical to the KDE plots for the training dataset (and have almost zero JSD measure).

### 5.1 Comparison with Markov Chain Monte Carlo Method on Synthetic Target Data

Although the results shown in Fig. 11B are encouraging, it is important to test the performance of the cGAN on data that were not part of the training dataset. To create synthetic target data to use for validation, we constructed 100 random parameter sets with each parameter *p* drawn from a normal distribution with mean *μ*_*p*_ and standard deviation *μ*_*p*_*/*8, where *μ*_*p*_ is the default value of that parameter. If the randomly drawn value was negative or was larger than the upper bound for that parameter in the training set, then the value was set to zero or the upper bound, respectively.

We simulated these 100 parameter sets with the CA1 model, calculated the features, and passed the features to the trained cGAN to generate 100 cGAN parameter samples. We then simulated these cGAN parameter samples with the CA1 model, calculated the features, and compared the target and cGAN feature distributions. KDE plots for each feature, as well as 2D KDE plots for each pair of features, show that the cGAN feature distributions are similar to the target feature distributions (Figs. 12 and 13A lower triangles). Furthermore, KDE plots for each parameter and each pair of parameters show that the cGAN parameter distributions are similar to the parameter distributions used to generate the target data (Fig. 13B lower triangle). To confirm these visual conclusions, we performed two-sample Kolmogorov-Smirnov (KS) tests to compare the cGAN and target distributions in feature and parameter space. In all cases but one (the voltage at the maximum positive rate of rise feature), the KS test failed to reject the null hypothesis (with a p-value threshold of 0.01) that these two sets of samples come from the same probability distribution, suggesting that the cGAN distributions are indeed similar to the target distributions.

**Fig. 12.**
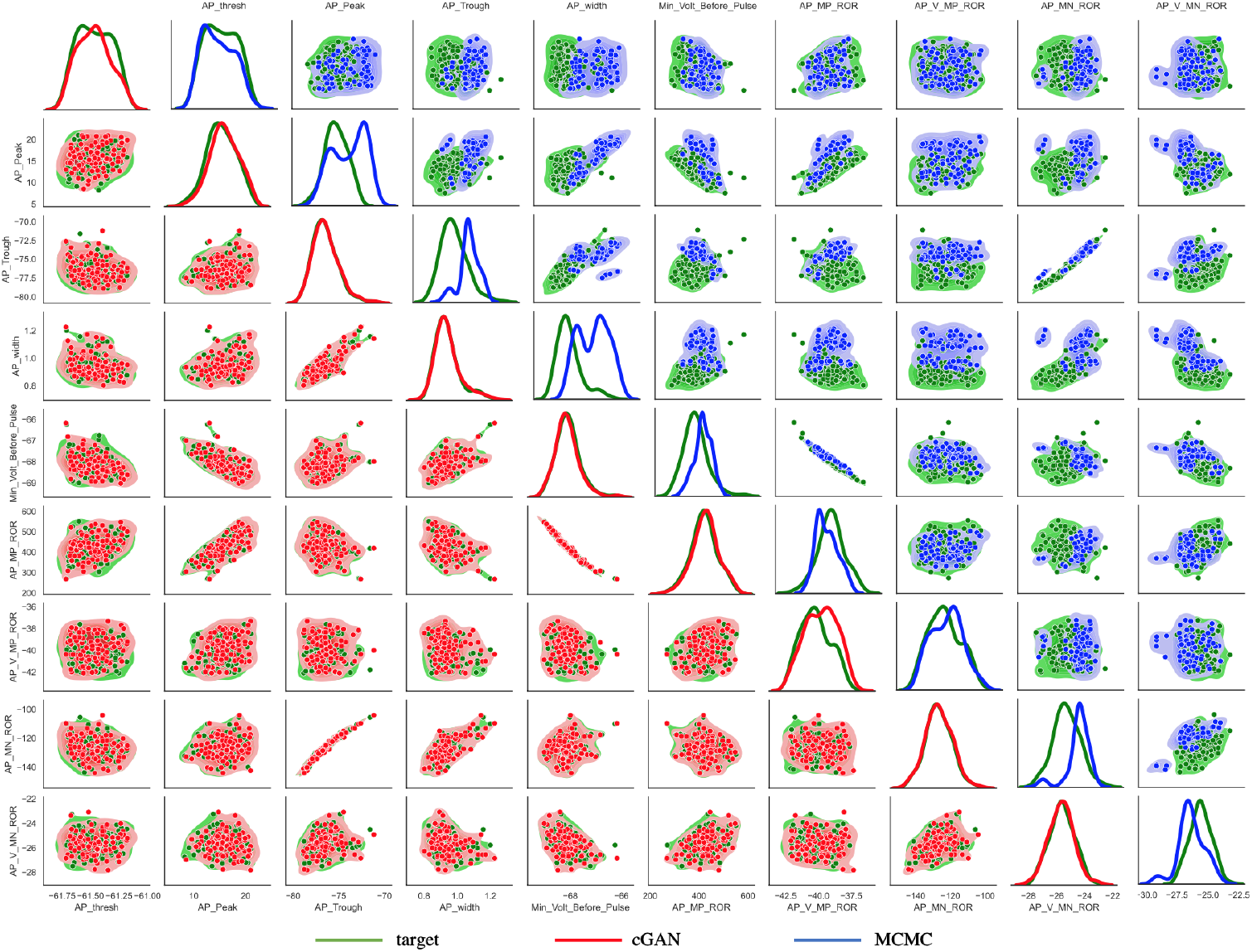
Performance of cGAN and MCMC on synthetic target data - AP features. *Lower main diagonal and lower triangle* - KDE and scatter plots of the cGAN samples (red) versus target (green). *Upper main diagonal and upper triangle* - KDE and scatter plots of the MCMC samples (blue) versus target (green).

**Fig. 13.**
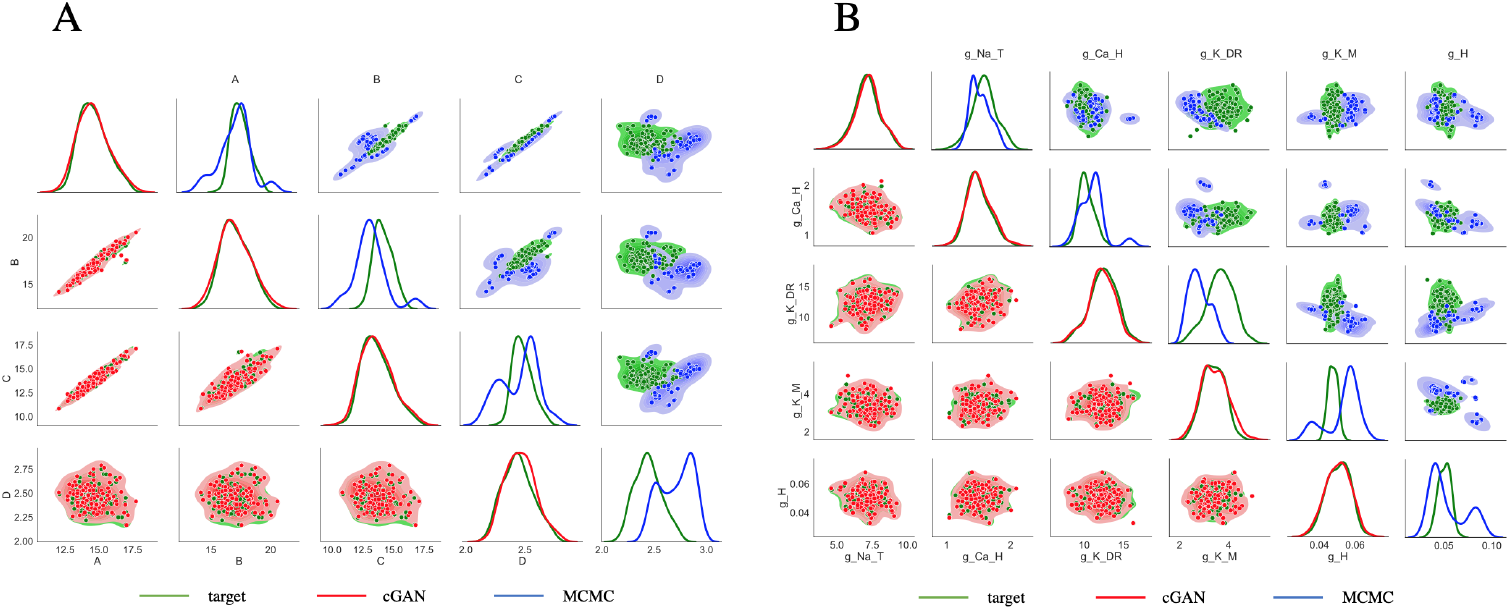
Performance of cGAN and MCMC on synthetic target data - HP features and parameter space. *Lower main diagonal and lower triangle* - KDE and scatter plots of the cGAN samples (red) versus target (green). *Upper main diagonal and upper triangle* - KDE and scatter plots of the MCMC samples (blue) versus target (green). **(A)** HP features. **(B)** Parameter space.

We then performed the same procedure using a Markov chain Monte Carlo (MCMC) method instead of cGAN to infer parameters from the target data. The details of our MCMC implementation are provided in the Supplementary Methods. We chose MCMC as the benchmark method to compare the performance of cGAN to because most state-of-the-art methods for solving stochastic inverse problems are based on MCMC [11, 33]. We passed the same 100 target data features to the MCMC algorithm as we did the cGAN, and then simulated the parameters produced by MCMC with the CA1 model. The feature distributions produced by the MCMC parameters, and the distributions of the MCMC parameters themselves, differ from their respective target distributions (Figs. 12 and 13 upper triangles). Furthermore, we performed KS tests between the MCMC parameters and features and the target parameters and features, and in all cases the null hypothesis that these samples come from the same probability distribution was rejected, suggesting that the MCMC distributions are indeed different from the target distributions.

### 5.2 Parameter Inference Tests on Synthetic Target Data

To further test the ability of cGAN to accurately infer biophysical model parameters from feature data, we generated synthetic target datasets with a variety of underlying parameter structures. These structures were chosen to reflect the possible scenarios one may encounter when working with data from 2 different categories (e.g. data from WT versus mutant mice, or data from young versus old mice). For the CA1 model, we are interested in 5 parameters. Suppose that in the mutant mice, only 1 of these parameters (e.g., *g*_*NaT*_) is altered compared to WT, and the other 4 parameters are not. To simulate this scenario, we construct two groups of target data. For each of the 4 parameters that are not hypothesized to be altered by the mutation (i.e., *g*_*CaH*_, *g*_*KDR*_, *g*_*KM*_, and *g*_*H*_), we draw 100 values from the same normal distribution 𝒩 (*μ*_*p*_, (*μ*_*p*_*/*8)^2^) for each group. For the parameter that is altered by the mutation (*g*_*NaT*_), we draw 100 values from 𝒩 (0.5*μ*_*p*_, (*μ*_*p*_*/*8)^2^) for Group 1 and 100 values from 𝒩 (1.5*μ*_*p*_, (*μ*_*p*_*/*8)^2^) for Group 2. For each group, we then: (1) simulate these parameter sets using the CA1 model and calculate the features, s(2) pass the features to the trained cGAN as target data to obtain cGAN samples (parameter sets), (3) simulate the cGAN parameter sets using the CA1 model and calculate the features, and (4) compare the cGAN feature and parameter distributions between Group 1 (G1) and Group 2 (G2) and to their respective targets. The G1 and G2 target distributions of some AP features are quite different from each other (Fig. S8 lower triangle), whereas the G1 and G2 membrane hyperpolarization feature target distributions are similar (Fig. S9 lower triangle), illustrating that the value of *g*_*NaT*_ affects some features more than others. Nonetheless, for all features the cGAN samples reproduce the target distributions well across both G1 and G2. Figure 14 (lower triangle) shows that the cGAN was able to accurately infer the distributions of all 5 parameters as well. Importantly, the cGAN-inferred distributions for *g*_*NaT*_ are distinct between Groups 1 and 2, whereas for the other 4 parameters the cGAN-inferred distributions are similar for G1 and G2.

**Fig. 14.**
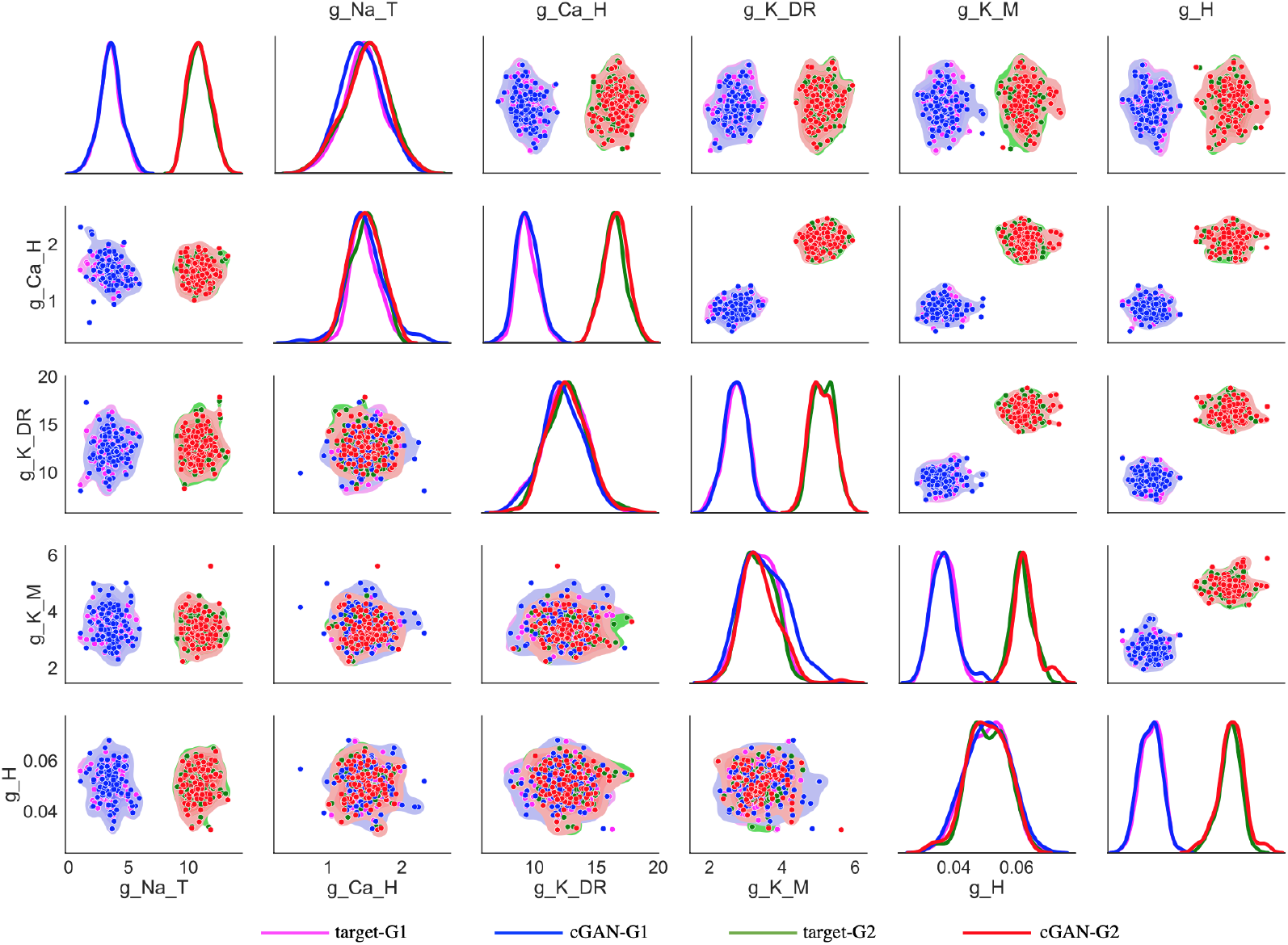
Performance of cGAN in parameter space on synthetic targets from 2 groups with distinct parameter structures. KDE plots (main diagonals) and scatter plots (lower and upper triangles) for Group 1 (G1) target data (magenta), Group 2 (G2) target data (green), cGAN samples for G1 (blue) and cGAN samples for G2 (red). *Lower main diagonal and lower triangle* - only 1 parameter (*g*_*NaT*_) is distributed differently in the G1 target data than in the G2 target data, and the other 4 parameters have the same distribution in the G1 and G2 target data. *Upper main diagonal and upper triangle* - Four parameters (all parameters except *g*_*NaT*_) are distributed differently in the G1 target data than in the G2 target data.

We also used KS tests to assess the cGAN performance. First, we ran KS tests on the target data from G1 and G2. For the parameters, the null hypothesis that the G1 and G2 target samples come from the same distribution is rejected for *g*_*NaT*_, but is not rejected for the other 4 parameters, as one would hope since this was the true structure used to create the target data. For the feature distributions, the KS tests reject the null for 10 out of the 13 features. When we ran the KS tests on the cGAN G1 and G2 distributions, we get the exact same results for both the parameters and features as we did for the target data. This consistency in KS test results suggests that cGAN is able to correctly identify the structure of parameter variations between two groups of samples based on their features. Additionally, we ran KS tests to compare the cGAN distributions for G1 to the target data for G1, and the cGAN distributions for G2 to the target data for G2. For G1, all of the KS tests (for both parameters and features) failed to reject the null, again indicating the cGAN distributions are similar to the target distributions. For G2, all of the KS tests failed to reject the null except for one feature (the voltage at the maximum positive rate of rise). We then repeated this simulation and testing procedure 4 more times with the other possible choices for having a single parameter distinguish G1 and G2. For the G1 vs. G2 KS tests, the cGAN sample tests returned the same result as the target data tests 70 out of 72 times (Supplementary Fig. S2 top panels). For the cGAN sample vs. target data KS tests, the null was rejected 0 out of 72 times for G1 (Fig. S2 bottom left panel) and 2 out of 72 times for G2 (Fig. S2 bottom right panel).

Next, we investigated scenarios with more than one parameter distinguishing G1 and G2. For example, we considered the case where *g*_*NaT*_ is not altered by the mutation, but the other 4 parameters all are altered by the mutation (i.e. *g*_*NaT*_ ∼ 𝒩(*μ*_*p*_, (*μ*_*p*_*/*8)^2^) in both G1 and G2, but *g*_*CaH*_, *g*_*KDr*_, *g*_*KM*_, and *g*_*H*_ are all distributed 𝒩 (0.5*μ*_*p*_, (*μ*_*p*_*/*8)^2^) in G1 and 𝒩 (1.5*μ*_*p*_, (*μ*_*p*_*/*8)^2^) in G2). The KDE and scatter plots for the AP features (Fig. S8 upper triangle), membrane hyperpolarization features (Fig. S9 upper triangle), and parameters (Fig. 14 upper triangle) indicate that the cGAN samples are consistent with the target data distributions. We also simulated the 4 other cases where each of the other 4 parameters was the only one not altered by the mutation. For the G1 vs. G2 KS tests, the cGAN sample tests returned the same result as the target data tests 89 out of 90 times (Fig. S5 top panels upper portion). For the cGAN sample vs. target data KS tests, the null was rejected 0 out of 90 times for G1 and 3 out of 90 times for G2 (Fig. S5 top panels, bottom left and bottom right portions, respectively).

There are 35 other ways that exactly 4 out of the 5 parameters could be altered by the mutation. In Fig. 14 (upper triangle), the other 4 parameters all had lower means in G1 than in G2. Instead, up to three of these parameters could have higher means in G1 than in G2 (if all 4 had higher means, it would be equivalent to the case we already simulated just with the G1 and G2 labels swapped). We simulated and tested these additional parameter structures. For the G1 vs. G2 KS tests, the cGAN sample tests returned the same result as the target data tests 624 out of 630 times (Fig. S5 bottom 7 panels upper portions). For the cGAN sample vs. target data KS tests, the null was rejected 15 out of 630 times for G1 and 38 out of 630 times for G2 (Fig. S5 bottom 7 panels lower portions).

So far, we have discussed scenarios where either exactly 1 parameter was affected by the mutation or exactly 4 parameters were affected by the mutation. Here, we consider the remaining scenarios of exactly 2, 3, or 5 parameters being affected. The number of possible cases for each scenario is given by

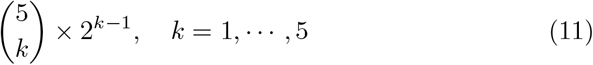

where *k* is the number of parameters affected by the mutation. Thus, for *k* = 2, we have 20 different cases. For the G1 vs. G2 KS tests, the cGAN sample tests returned the same result as the target data tests 353 out of 360 times (Fig. S3 upper portions of panels). For the cGAN sample vs. target data KS tests, the null was rejected 3 out of 360 times for G1 and 15 out of 360 times for G2 (Fig. S3 lower portions of panels). For *k* = 3, there are 40 different cases. For the G1 vs. G2 KS tests, the cGAN sample tests returned the same result as the target data tests 711 out of 720 times (Fig. S4 upper portions). For the cGAN sample vs. target data KS tests, the null was rejected 9 out of 720 times for G1 and 38 out of 720 times for G2 (Fig. S4 lower portions). Finally, for *k* = 5, we have 16 different cases. For the G1 vs. G2 KS tests, the cGAN sample tests returned the same result as the target data tests 288 out of 288 times (Fig. S6 upper portions). For the cGAN sample vs. target data KS tests, the null was rejected 9 out of 288 times for G1 and 13 out of 288 times for G2 (Fig. S6 lower portions). The results of the KS tests for all of the 5 choose *k* cases are summarized in Fig. S7.

In summary, these results on synthetic target data demonstrate that cGAN is capable of accurately identifying complex parameter variation structures from subtle differences in the features of CA1 model simulations. This gives us the confidence to apply the cGAN method to experimental data in Section 6.

## 6 Parameter Inference on Experimental Target Data

Now that we have established cGAN as a tool for mapping observed traces to unobserved/unmeasured parameter values, we turn our attention back to experimental data (Figs. 1, 3, 4) and seek to answer the following questions: Which maximal conductances are responsible for the differences observed in feature space between (1) the wild type versus the mutant mice (i.e. the disease effect), and (2) the 12-month old mice versus the 24-month old mice (i.e. the age effect)?

To answer these questions, we will pass the features for each cell to the cGAN to obtain a population of models for each individual cell. For some cells, certain feature values fall outside the range of the values for that feature used in our training dataset. This can lead to inaccurate cGAN samples; thus, if a cell has a feature value outside that range we replaced that value with the median value of that feature across the training dataset. We obtained 100 cGAN samples for each cell, and then pushed those parameters forward through the mechanistic model. Figure 15A shows that the mean AP and hyperpolarization traces produced by the cGAN samples agree well with the mean AP and hyperpolarization traces from the experimental recordings in each of the 4 categories. For example, we can see from the voltage traces that the AP peak feature exhibits the same trend in the cGAN samples and experimental data, with WT 24 month having the highest mean AP peak, followed by PDAPP, WT 12 month, and rTG4510, respectively. To give a sense of how the variability of the cGAN samples compares to the experimental recordings, Fig. 15B shows the mean *±* standard deviation of the AP and hyperpolarization traces. Overall, the amount of variability in the cGAN samples appears comparable to the amount of variability in the experimental data across the 4 categories, with the exception of the hyperpolarization traces for WT 24 month where there is less variability in the cGAN samples than in the data. Furthermore, box-and-whisker plots for the output of the cGAN samples in feature space also agree well with the feature distributions in the experimental data (compare Fig. S10 to Fig. 3, and Fig. S11 to Fig. 4).

**Fig. 15.**
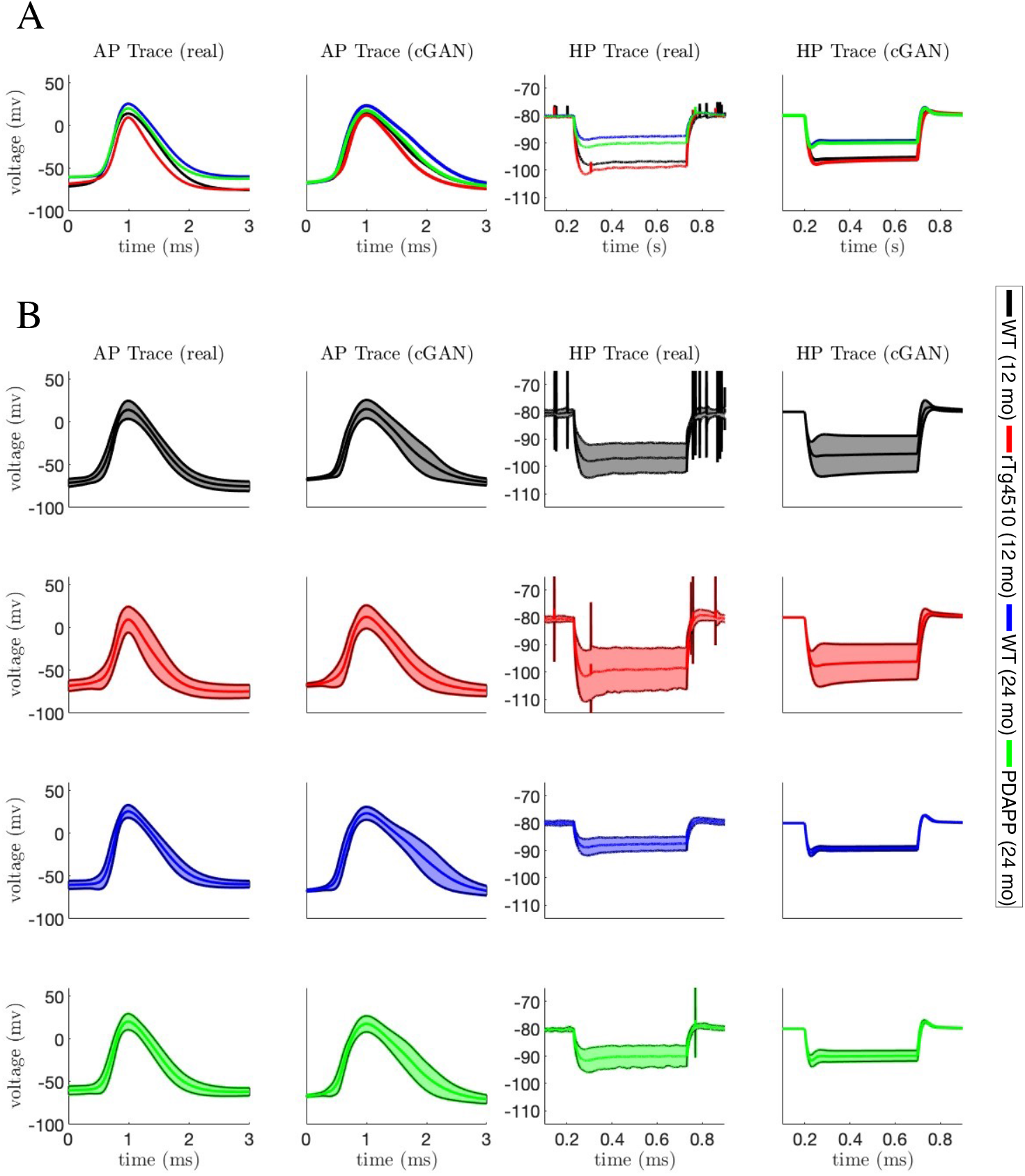
AP and membrane hyperpolarization traces from cGAN samples with experimental targets. **(A)** Mean AP and membrane hyperpolarization traces from each experimental data category (1st and 3rd panels) and mean AP and membrane hyperpolarization traces from simulated the mechanistic model with 100 cGAN parameter samples for each cell in each category (2nd and 4th panels). **(B)** Same as A1, but shading shows the mean *±* standard deviation for each category.

Seeing that the cGAN samples produce appropriate behavior in feature space, we move on to assessing the distributions of the cGAN samples in parameter space in order to answer the questions posed at the beginning of this section. First, we compare the cGAN samples for rTg4510 and their age-matched controls (WT 12 month). Based on KDE plots for each parameter (lower main diagonal of Fig. 16), we see that for 3 of the parameters (*g*_*NaT*_, *g*_*CaH*_, *g*_*H*_), the WT 12 month and rTg4510 distributions are centered around the same values. However, the distribution for *g*_*KDR*_ is shifted to the right in rTg4510 relative to WT 12 month, whereas the distribution of *g*_*KM*_ is shifted to the left in rTg4510 relative to WT 12 month. The KDE plots comparing PDAPP and their age-matched controls (WT 24 month) show a similar trend, with the *g*_*KDR*_ and *g*_*KM*_ distributions shifted to the right and left, respectively, for PDAPP relative to WT (upper main diagonal of Fig. 16). For PDAPP, the distribution of *g*_*NaT*_ is also shifted to the left relative to WT. From these observations, we hypothesize that for the mouse model of tauopathy, it is the conductances *g*_*KDR*_ and *g*_*KM*_ that are responsible for the altered excitability properties. For the mouse model of amyloidopathy, we hypothesize that these conductances plus *g*_*NaT*_ play a role in the altered excitability.

**Fig. 16.**
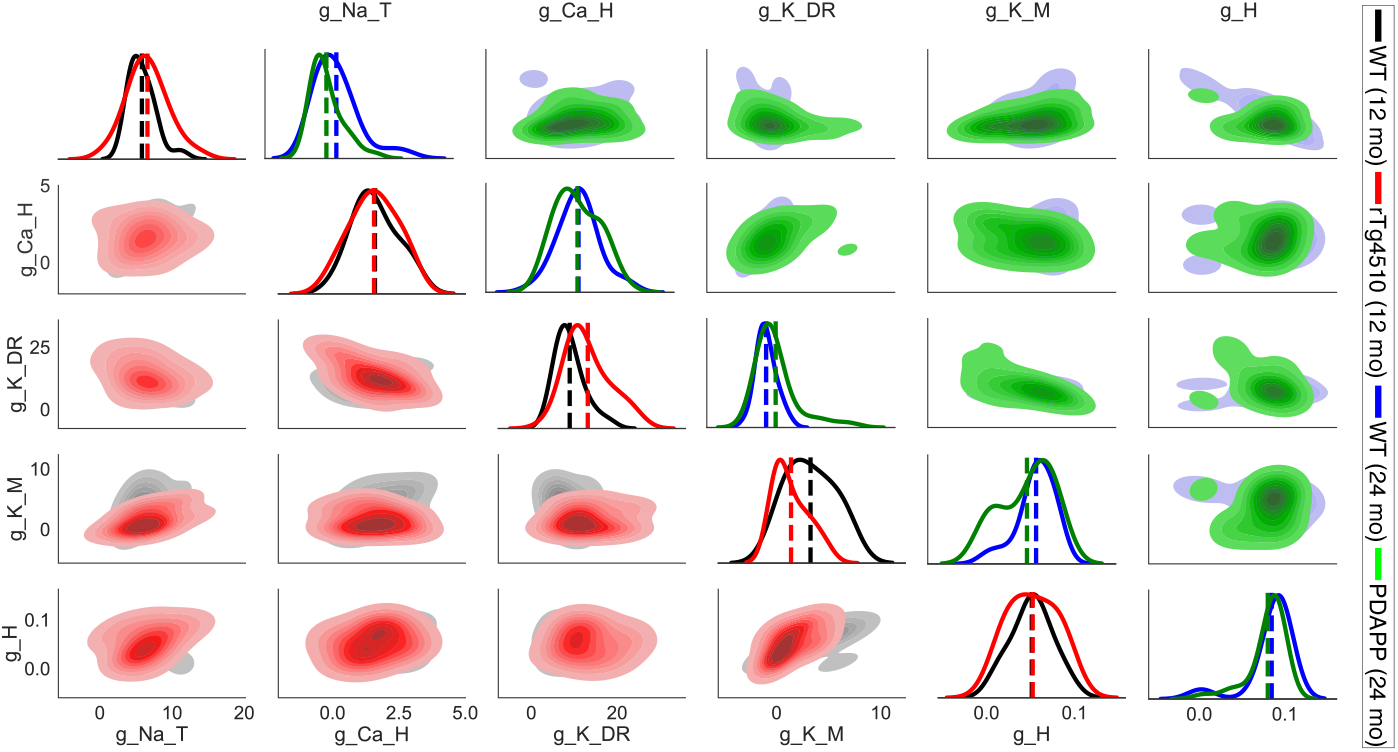
Disease effect - parameter distributions from cGAN samples with experimental targets. *Lower main diagonal and lower triangle* - KDE and shaded contour plots of cGAN samples for 12-month-old WT (black) and 12-month-old tau mutant (rTg4510, red) mice. *Upper main diagonal and upper triangle* - KDE and shaded contour plots of cGAN samples for 24-month-old (blue) and 24-month-old amyloid beta mutant (PDAPP, green) mice.

Having considered the disease effect, we now move on to assessing the age effect. First, we compare the cGAN samples for WT 12 month and WT 24 month. Based on KDE plots for each parameter (lower main diagonal of Fig. 17), we see the most striking differences for *g*_*H*_, with the distribution for the older mice shifted to the right relative to the distribution for the younger mice. The parameter *g*_*KM*_ also shows a rightward shift in the older mice. On the other hand, the distribution for *g*_*KDR*_ is shifted to the left in the older mice. The 12 to 24 month WT comparison is the most appropriate one for assessing an age effect, since the only difference between these two groups of mice is their age. However, for the sake of completeness we also compared the 12 month mutant (rTg4510) to the 24 month mutant (PDAPP). Remarkably, the 3 shifts in the parameter distributions that we observed with age in the WT mice were all preserved in the mutant mice, despite the fact that the 12 and 24 month mutants have different mutations (tauopathy and amyloidopathy, respectively). Specifically, the *g*_*H*_ and *g*_*KM*_ distributions are shifted to the right in the older mutants relative to the younger mutants, while *g*_*KDR*_ is shifted to the left in the older mutants (upper main diagonal of Fig. 17). These results lead us to hypothesize that these 3 conductances underlie the changes in excitability properties observed with aging.

**Fig. 17.**
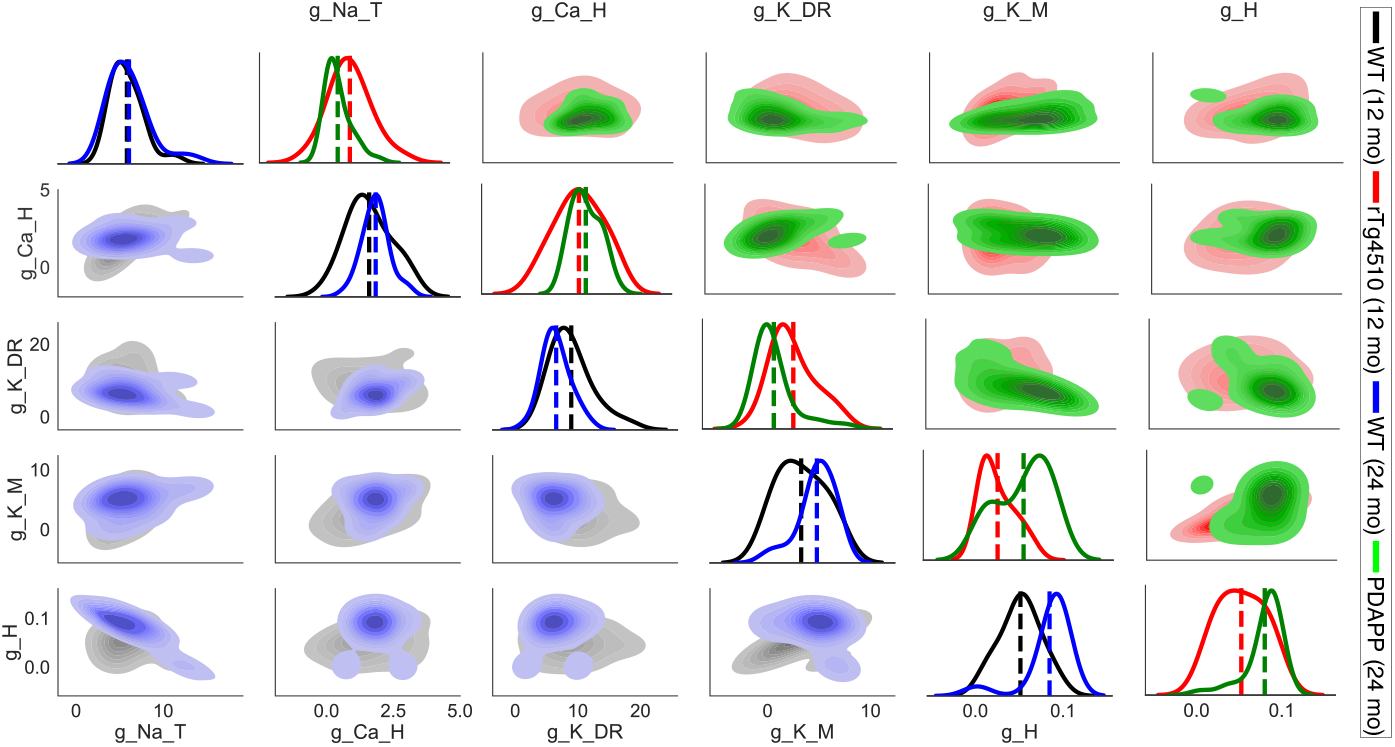
Age effect - parameter distributions from cGAN samples with experimental targets. *Lower main diagonal and lower triangle* - KDE and shaded contour plots of cGAN samples for 12-month-old (black) and 24-month-old (blue) WT mice. *Upper main diagonal and upper triangle* - KDE and shaded contour plots of cGAN samples for 12-month-old (rTg4510, red) and 24-month-old (PDAPP, green) mutant mice.

## 7 Discussion

The last decade has seen a rise in the application of population-based modeling in the neuronal and cardiac electrophysiology domains [13–15, 34–37]. The development of methods for selecting and producing virtual patient populations that accurately reflect the statistics of clinical populations has also received a lot of attention in fields such as quantitative systems pharmacology [16, 18, 19, 38, 39].

Here, we have introduced and employed a deep hybrid modeling (DeepHM) framework [17, 20] featuring conditional generative adversarial networks (cGANs) that can be categorized as a population of models technique. We compared the performance of cGAN and a standard Bayesian inference Markov chain Monte Carlo (MCMC) method [40] on a parameter inference task with synthetic target data where the ground truth was known. The cGAN outperformed MCMC on this task, and showed it is capable of producing a population of models that captures the type of variability that is often present in biological data. Since the cGAN was able to accurately detect a variety of complex differences in the distributions of parameters across 2 groups of synthetic target data, we employed the cGAN to infer the parameter distributions across 4 groups of experimental target data (WT and mutant mice at 2 different ages). From these distributions, we drew conclusions about the biophysical mechanisms (i.e. the ionic conductances) underlying the differences in the observed excitability properties of WT versus mutant and younger versus older mice. These results illustrate the value of mapping experimental data back to the parameter space of a mechanistic model. In future work, the predictions we made about the role of certain conductances in Alzheimer’s disease and aging phenotypes can be tested experimentally.

Since DeepHM can produce populations of cell models that accurately reflect the heterogeneous responses of real cells, this framework could prove useful in virtual drug design applications. This future direction is inspired by recent work where a population of models approach was used to identify a set of ion channel drug targets that optimally convert Huntington’s disease cellular phenotypes to healthy phenotypes simultaneously across multiple measures [41].

In this paper, we considered a stochastic inverse problem (SIP), where the experimental data comes from multiple individuals across a population. Our method utilized recent advances in deep learning to generate model parameter sets that produced a population of deterministic mechanistic models with outputs that are consistent with the experimental population data. A distinct but related problem is simulation-based inference (SBI), where experimental data are acquired from a single individual and a stochastic mechanistic model is used to infer the set of model parameters most likely to have generated the data distribution observed from the individual. While deep learning methods such as neural density estimation with normalizing flows have been used in SBI problems [9, 42], to our knowledge our work is among the first to apply a deep learning approach to an SIP.

## Acknowledgments

This work was partially supported by NSF DMS grants 1555237 and 2152115 to COD. KCAW was graciously supported by the EPSRC New Horizons grant (EP/V048716/1) and the EPSRC Hub for Quantitative Modeling in Healthcare (EP/T017856/1). The PDAPP data were collected under the Medical Research Council grant G1100623 (FT), and the rTg4510 data under the Alzheimer’s Society Junior Fellowship grant AS-JF-14-007 (FT). KCAW and FT received support from the the Wellcome Trust ISSF2 Centre grant WT105618MA.

## Data Availability Statement

The datasets generated during and/or analysed during the current study are available from the corresponding author on reasonable request.

## Notes

### Competing Interest Statement

The authors have declared no competing interest.

