## Supplementary Information for "Inferring parameters of pyramidal neuron excitability in mouse models of Alzheimer’s disease using biophysical modeling and deep learning"

### 1 Mechanistic Model Parameters

**Table 1 Parameters of CA1 pyramidal neuron model.** For most parameters, we used the values given in Nowacki et al. [1] paper. Parameter values for the hyperpolarization-activated potassium current ($I_H$) are taken from Booth et al. [2]. Optimized parameter values for the maximal conductances and transient sodium half-activation that we obtained through differential evolution (DE-MG-Vmnat) are shown in parentheses.

| Parameter | Value | Units | Parameter | Value | Units |
|---|---|---|---|---|---|
| $C_m$ | 1 | $\mu$F/cm$^2$ | $V_{m_H}$ | -102 | mV |
| $E_{Na}$ | 60 | mV | $V_{n_H}$ | -102 | mV |
| $E_{Ca}$ | 90 | mV | $k_{m_{Na_T}}$ | 5 | mV |
| $E_K$ | -85 | mV | $k_{h_{Na_T}}$ | -7 | mV |
| $E_H$ | -30 | mV | $k_{m_{Na_P}}$ | 3 | mV |
| $E_L$ | -65 | mV | $k_{m_{Ca_T}}$ | 5 | mV |
| $g_{Na_T}$ | 65.0 (7.2603) | $\mu$S/cm$^2$ | $k_{h_{Ca_T}}$ | -8.5 | mV |
| $g_{Na_P}$ | 0.1 (0.0423) | $\mu$S/cm$^2$ | $k_{m_{Ca_H}}$ | 5 | mV |
| $g_{Ca_T}$ | 0.6 (0.067) | $\mu$S/cm$^2$ | $k_{h_{Ca_H}}$ | -7 | mV |
| $g_{Ca_H}$ | 0.74 (1.5208) | $\mu$S/cm$^2$ | $k_{m_{K_{DR}}}$ | 11.4 | mV |
| $g_{K_{DR}}$ | 9.5 (12.505) | $\mu$S/cm$^2$ | $k_{h_{K_{DR}}}$ | -9.7 | mV |
| $g_{K_M}$ | 0.8 (3.3837) | $\mu$S/cm$^2$ | $k_{m_{K_M}}$ | 10 | mV |
| $g_H$ | 0.05 (0.0503) | $\mu$S/cm$^2$ | $k_{m_H}$ | -13 | mV |
| $g_L$ | 0.02 (0.0035) | $\mu$S/cm$^2$ | $k_{n_H}$ | -6 | mV |
| $V_{m_{Na_T}}$ | -37 (-60.00) | mV | $\tau_{m_{Ca_T}}$ | 2 | ms |
| $V_{h_{Na_T}}$ | -75 | mV | $\tau_{h_{Ca_T}}$ | 32 | ms |
| $V_{m_{Na_P}}$ | -47 | mV | $\tau_{m_{Ca_H}}$ | 0.08 | ms |
| $V_{m_{Ca_T}}$ | -54 | mV | $\tau_{h_{Ca_H}}$ | 300 | ms |
| $V_{h_{Ca_T}}$ | -65 | mV | $\tau_{m_{K_{DR}}}$ | 1 | ms |
| $V_{m_{Ca_H}}$ | -15 | mV | $\tau_{h_{K_{DR}}}$ | 1400 | ms |
| $V_{h_{Ca_H}}$ | -60 | mV | $\tau_{m_{K_M}}$ | 75 | ms |
| $V_{m_{K_{DR}}}$ | -5.8 | mV | $\tau_{m_H}$ | 15 | ms |
| $V_{h_{K_{DR}}}$ | -68 | mV | $\tau_{n_H}$ | 210 | ms |
| $V_{m_{KM}}$ | -30 | mV | $p$ | 0.85 | - |

### 2 Supplementary Methods

#### 2.1 Differential Evolution (DE) - global optimization

Differential evolution is a type of stochastic global optimization and a population-based search technique first introduced by Storn and Price, 1997 [3, 4]. This method is comprised of four different steps, including *Initialization*, *Mutation*, *Crossover*, and *Selection.*

*Initialization* - a population of individuals, which is also known as the first generation or parents, is often defined by drawing a population number randomly from a uniform distribution to explore the objective landscape. In this context an individual is essentially an ordered set of parameters.

*Mutation* - a new vector (a.k.a donor vector) is introduced in this step by a linear combination of three random vectors from the current generation but not the one which we are going to compare and substitute with in the next generation (i.e. target vector), this means at least four members are needed for applying this method.

*Crossover* - the recombination of donor vector with the best member of the previous generation with a probability rate bring both exploration and exploitation to the search and lead the system get out of the local minima. *Selection* - during the selection step, we have a new vector, which is a candidate for going to the new generation in replace of the target vector, if it satisfies certain conditions, and it has a better score [1] than the target vector. In this way, after each generation we might have better members or in the worst-case scenario all members could be the same as the previous generation without any change but none of them getting worse than before.

The Differential Evolution Algorithm that we used in this project known as **classic DE** in [3]. It is also referred to as DE/rand/1/bin which means the base vector is *randomly* selected and only *1* vector difference is added in the mutation part. Finally, the donated parameters in the crossover section follow a *binomial* distribution.

#### 2.2 Sobol indices - global sensitivity analysis

Sensitivity Analysis (SA) [5–9] addresses the question of how much the output of a model changes given a change in the input. There are two main classes of sensitivity analysis, *local* and *global.* In local SA, the goal is to deal with sensitivity in the neighborhood of a particular parameter value. Taking a derivative is an example of local sensitivity analysis as it only consider the behavior of the system

[1] The score could be the output of the objective function and depends on the minimization or maximization problem. The new candidate will be either rejected or accepted for the next generation.

near a single point. Global SA, on the other hand, focuses on the variability of model outputs throughout parameter space. Global SA provides more information about the system and because of that it is often preferred, but if the system is too large the computational cost of this technique can be prohibitive. There are several different global SA approaches, including linear methods, tree-based methods, regionalized sensitivity analysis (also known as Monte Carlo filtering methods), and variance-based techniques [10, 11]. Sobol SA is a variance-based technique that considers the contribution of the input parameters to the variance of the system outputs. In Sobol SA we typically use two measures, the first order index and total effect index. The first order index is the contribution of an individual parameter to the response variance without considering any interactions with the other parameters. The total effect index is the contribution of an individual parameter to the response variance that does consider interactions with the other parameters.

Suppose we have a function of $f$ that maps the vector of variables $X = (X_1, X_2, \cdots, X_p)$ to some quantity of interest. We assume $X$ has some known probability distribution, this correspond to certain parameters in the model which this probability distribution reflect the level of uncertainty in them. The goal is to determine the sensitivity of $f$ to $X$, under the assumption that $f$ is square integrable.

$$\begin{aligned} f\colon \mathbb{R}^p &\to \mathbb{R} \\ X &\mapsto f(X) \end{aligned}$$

If $f$ be square integrable (i.e. $f(x) \in L^2$), we can decompose the above function as the sum of the functions of only $1, 2, \cdots, p$ parameters:

$$f(X) = f_0 + \sum_{i=1}^{p} f_i(X_i) + \sum_{1 \le i < j \le p} f_{i,j}(X_i, X_j) + \cdots + f_{1,2,\cdots,p}(X_1, X_2, \cdots, X_p) \tag{1}$$

where:

$$
\begin{aligned}
f_0 &= \mathbb{E}[f(X)] \\
f_i(X_i) &= \mathbb{E}[f(X)\|X_i] - f_0 \\
f_{i,j}(X_i, X_j) &= \mathbb{E}[f(X)\|X_i, X_j] - f_i(X_i) - f_j(X_j) - f_0
\end{aligned}
$$

If $X_1, X_2, \cdots, X_p$ are statistically independent then all $f$ satisfying the orthogonality property, which includes:

$$
\begin{aligned}
Var(f(X)) &= \sum_{k=1}^{p} D_k(X_k) + \sum_{1 \leq k \leq k' \leq p} D_{k,k'}(X_k, X_{k'}) + \cdots + D_{1,2,\cdots,p} \\
&= \sum_{u} D_u(X_u), \quad u \subset \{1, 2, \cdots, p\}
\end{aligned}
$$

where:

$$
D_k(X_k) = Var\left[f_k(X_k)\right], \quad D_{k,k'}(X_k, X_{k'}) = Var\left[f_{k,k'}(X_k, X_{k'})\right]
$$

$$
D_0 = Var\left[f_0\right] = 0, \quad \text{and} \quad u = \{i_1, i_2, \cdots, i_s\}, \quad 1 \leq s \leq p
$$

This means the total variance of the response $f(X)$ can be written as the sum of partial variances. By defining the total variance of the response $f(X)$ that can be attributed to input parameter $X_k$ as the ratio of $Var\left[\mathbb{E}[f(X)\|X_k]\right]/Var(f(X))$, the Sobol index for a subset **u** (i.e. $u \subset \{1, \cdots, p\}$) can be defined as the ratio between the contribution given by the interaction among the components of **u** for the model variance, and the total variance itself. Thus, the Sobol index for a subset **u** can be written as:

$$
S_u = \frac{D_u(X_u)}{Var(f(X))}, \quad \sum_{u \subset \{1,\cdots,p\}} S_u = \frac{\sum_u D_u(X_u)}{Var(f(X))} = \frac{Var(f(X))}{Var(f(X))} = 1 \tag{2}
$$

As was mentioned before, in Sobol SA, two measures (i.e. the first order index and the total index) are usually computed. The first order index refers to the contribution of any one parameter to the output variance and can be defined as:

$$S_i = \frac{D_i(X_i)}{Var(f(X))}, \quad i = 1, \cdots, p.$$

The total index refers to the contribution of all subsets with more than one parameter to the output variance, which means:

$$\begin{aligned} S_T &= \sum_{1 \leq i \leq j \leq p} S_{i,j} + \cdots + S_{1,\cdots,p} = \sum_{u \subset \{1,\cdots,p\}} S_u \\ &= 1 - S_i, \quad i \in u. \end{aligned}$$

The first order index (i.e. $S_i$) describes the impact of $X_i$ individually on the defined output, whereas, the total index (i.e. $S_T$) represents the effect of $X_i$ along with the interaction of other input variables. A high value for either of these two indices suggests that $X_i$, either alone or in conjunction with other input variables, has a considerable overall impact on the output space.

##### 2.3 Markov Chain Monte Carlo - MCMC

The ***Monte Carlo*** method [12, 13] is a stochastic technique that aims to calculate numerical results from many random samples. In other words, if a method uses random numbers to solve a problem, that method is a type of Monte Carlo method. By employing this randomness, it is possible that one can address some problems that, in theory, may have deterministic solutions that are hard to obtain. In order to draw some samples from a known distribution, this method repeatedly produces some random samples coming from that distribution. By repeating this process, eventually in the long run all the samples come from the desired distribution. One of the main issues with using a Monte Carlo method is that is a memoryless process, which can lead to a very long and time consuming process to draw samples from a complex distribution. In other words, if one sample comes from a high density region of the distribution, there is no guarantee that the next candidate will also come from the same highly probable region. A ***Markov Chain*** is a sequence of events where the probabilities of the future depend only on the present [14, 15]. By incorporating a Markov process into the Monte Carlo method, which is known as ***Markov Chain Monte Carlo (MCMC)***, information about the previous step affects the next proposal candidate, and the process of sampling from complex distributions is sped up. The Metropolis Hastings algorithm (Algorithm 1) is a particular MCMC method that performs a random walk through the probability distribution to try to evaluate the values with higher probability. The main idea behind Metropolis-Hastings is that by running this algorithm for a good amount of iterations, the random walk procedure helps the Monte Carlo method to draw samples from the stationary distribution, which is the posterior distribution or the target distribution. The important fact here is that based on this algorithm, the acceptance probability that comes from the density of the model output is computed using a Gaussian mixture model that is fit to the target dataset.

**Algorithm 1** Markov Chain Monte Carlo method - Metropolis Hastings

Define $f(x)$ as the target distribution, $N$ as the total number of iterations, $x_i$ is the current value and $q(x\|x_i)$ be a proposal distribution. This proposal distribution can be symmetric or asymmetric in this method. Define $x_0$ randomly from its defined range (i.e. $x_0 = x_l + rand(0,1) \times (x_u - x_l)$).

**while** Iter $<$ $N$ **do**
  $x^\star \sim q(x\|x_i)$, proposed candidate
  $\rho = min\left\{1, \frac{f(x^\star)q(x_i\|x^\star)}{f(x_i)q(x^\star\|x_i)}\right\}$, $u \sim U(0,1)$,
  **if** $u < \rho$
    $x_{i+1} = x^\star$
  **else**
    $x_{i+1} = x_i$
  **end**
  Iter = Iter + 1
**end**

#### 2.4 Jensen Shannon Divergence

$$\mathrm{JSD}(p\|q) = \frac{1}{2}\left\{\int p(x)\log\left(\frac{p(x)}{M(x)}\right)dx + \int q(x)\log\left(\frac{q(x)}{M(x)}\right)dx\right\},$$

where

$$M = \frac{p+q}{2}.$$

### 3 Supplementary Results

#### 3.1 Synthetic target methodology

As we are dealing with 5 parameters in our mechanistic model, we design different target data scenarios with 5 choose $k$ parameters distinguishing two different groups of target data.

***5 choose 0:*** In this scenario, since $k = 0$, we only have one target group. All 5 parameters are drawn from a normal distribution with a mean $\mu$ and standard deviation $\mu/8$, where $\mu$ is the value of that parameter in the optimized DE model parameter set (DE-MG-Vmnat). There is only one case to consider in this scenario.

***5 choose 1:*** In this scenario, since $k = 1$, we choose one of the 5 parameters to draw from a different distribution for the Group 1 (G1) and Group 2 (G2) target datasets. For that parameter, we draw the G1 samples from a normal distribution with a mean of $0.5\mu$ and the G2 samples from a normal distribution with a mean of $1.5\mu$. For both groups the normal distribution has a standard deviation of $\mu/8$, where again $\mu$ is the value of that parameter in the optimized DE model. There are 5 different cases in this scenario, since there are 5 parameters that can be chosen to distinguish G1 and G2.

***5 choose*** $k$***:*** Here we consider the scenarios $k \in \{2, 3, 4, 5\}$. The number of cases for each value of $k$ can be computed from Eqn. (3). The $2^{k-1}$ term reflects the number of different ways $k$ parameters can be different between G1 and G2. For example, suppose $k = 3$, and the 3 parameters chosen to be distributed differently between G1 and G2 are *g-Na-T*, *g-Ca-H*, and *g-K-DR*. There are $2^2$ different possibilities: (1) all 3 parameters are low in G1 and high in G2 – denoted L-H, L-H, L-H; (2) *g-Na-T* is high in G1 and *g-Ca-H*, *g-K-DR* are low in G1 – denoted H-L, L-H, L-H; (3) *g-Na-T* low in G1, *g-Ca-H* high in G1, and *g-K-DR* low in G1 – denoted L-H, H-L, L-H; and (4) *g-Na-T*, *g-Ca-H* low in G1 and *g-K-DR* high in G1 – denoted L-H, L-H, H-L. We do not have to simulate possibilities such as H-L, H-L, H-L or L-H, H-L, H-L, since they are equivalent to possibilities (1) and (2) listed above, respectively, just with the labels swapped for G1 and G2 swapped.

$$\binom{n}{k} \times 2^{k-1}, \quad k = 1, \cdots, 5 \tag{3}$$

#### 3.2 Kolmogorov Smirnov tests

We performed Kolmogorov Smirnov tests (KS-tests) to compare the cGAN samples to the ground truth target samples for all the different 5 choose $k$ synthetic target scenarios. The null hypothesis for these tests is that the two samples are from the same probability distribution. The plots in Figs. S1-S6 represent the results of these tests, where black indicates the null hypothesis is rejected (p-value $\leq$ 0.01) and peach indicates the null hypothesis is not rejected (p-value $>$ 0.01).

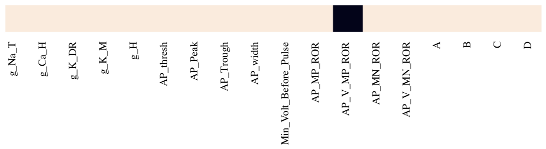


**Fig. S1 5 choose 0 - KS tests for cGAN samples versus target data samples.** Peach color indicates failure to reject the null hypothesis that the cGAN samples and synthetic target data samples are from the same distribution.

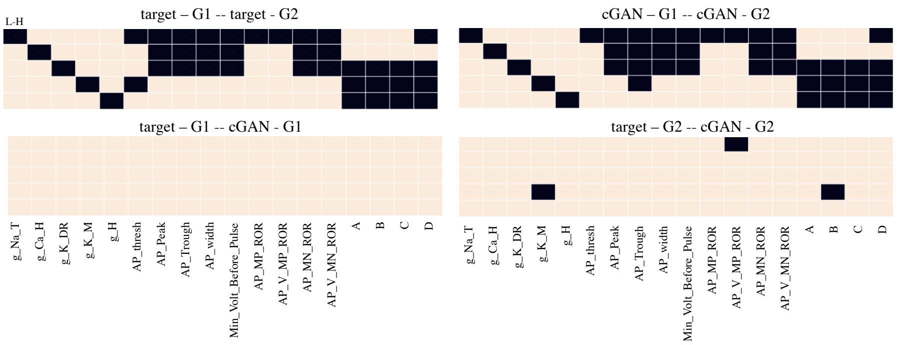


**Fig. S2 5 choose 1 - KS tests.** ***Top left*** - KS tests on two groups of targets (i.e. G1 (G2) with low - **L** (high - **H**) values of the parameter). ***Top right*** - KS test on cGAN samples corresponding to the two groups of target data (i.e. cGAN-G1 (cGAN-G2) with low - **L** (high - **H**) values of the parameter). ***Bottom left*** - KS test of cGAN-G1 versus target-G1. ***Bottom right*** - KS test of cGAN-G2 versus target-G2.

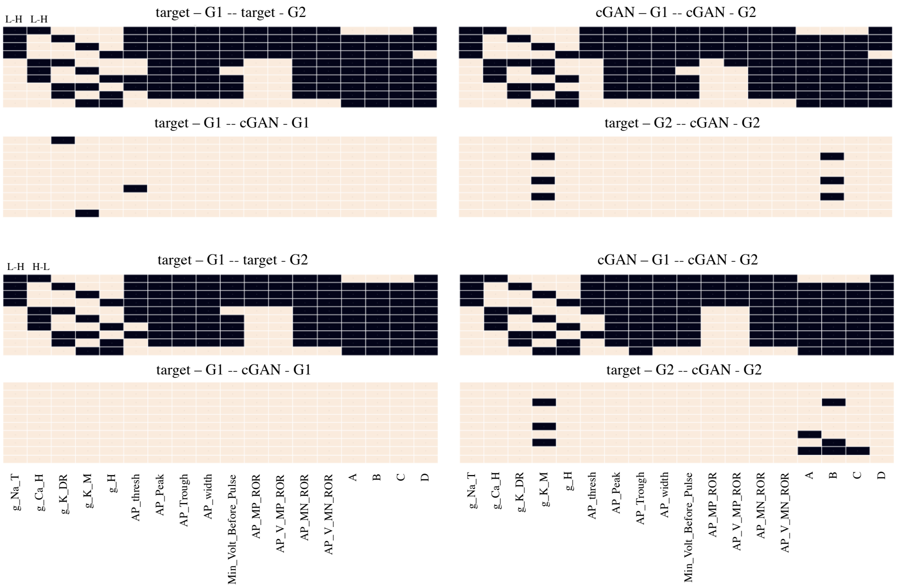


**Fig. S3 5 choose 2 - KS tests.** Panels are arranged in a similar fasion as Fig. S2. Top: L-H, L-H. Bottom: L-H, H-L.

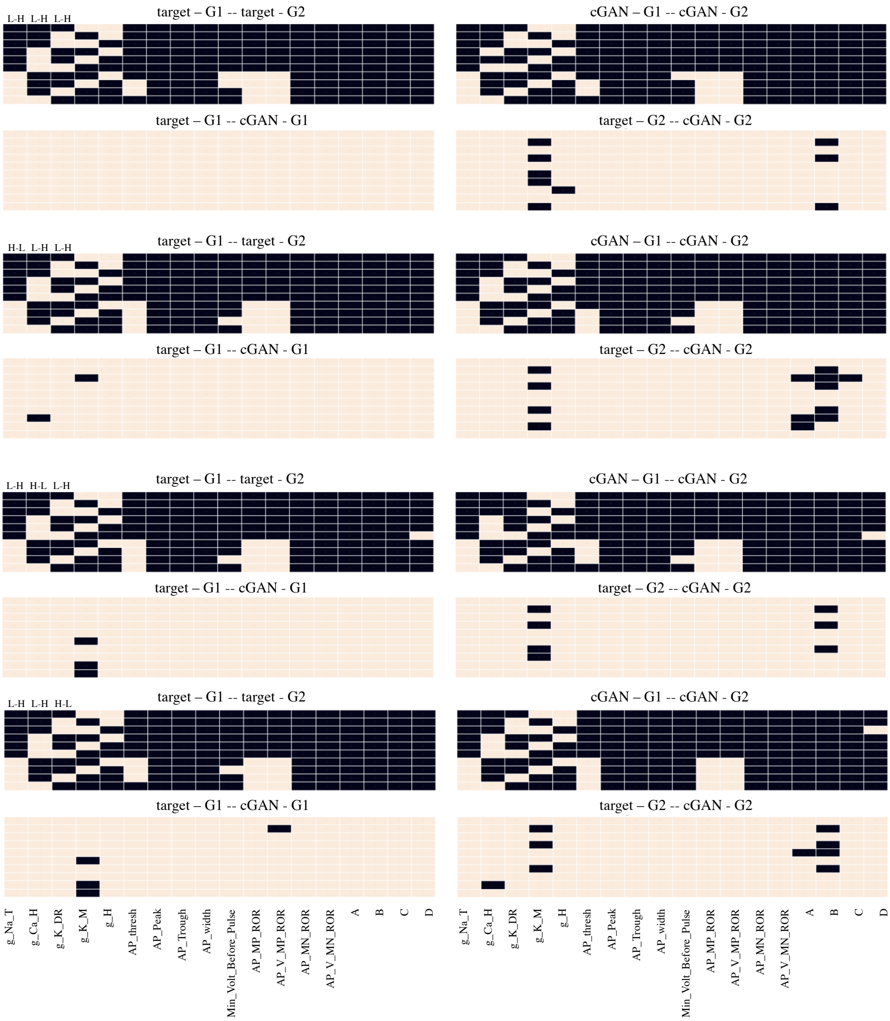


**Fig. S4 5 choose 3 - KS tests.** Panels are arranged in a similar fashion as Fig. S2. From top to bottom: ***(1) L-H L-H L-H***, ***(2) H-L L-H L-H***, ***(3) L-H H-L L-H***, ***(4) L-H L-H H-L***.

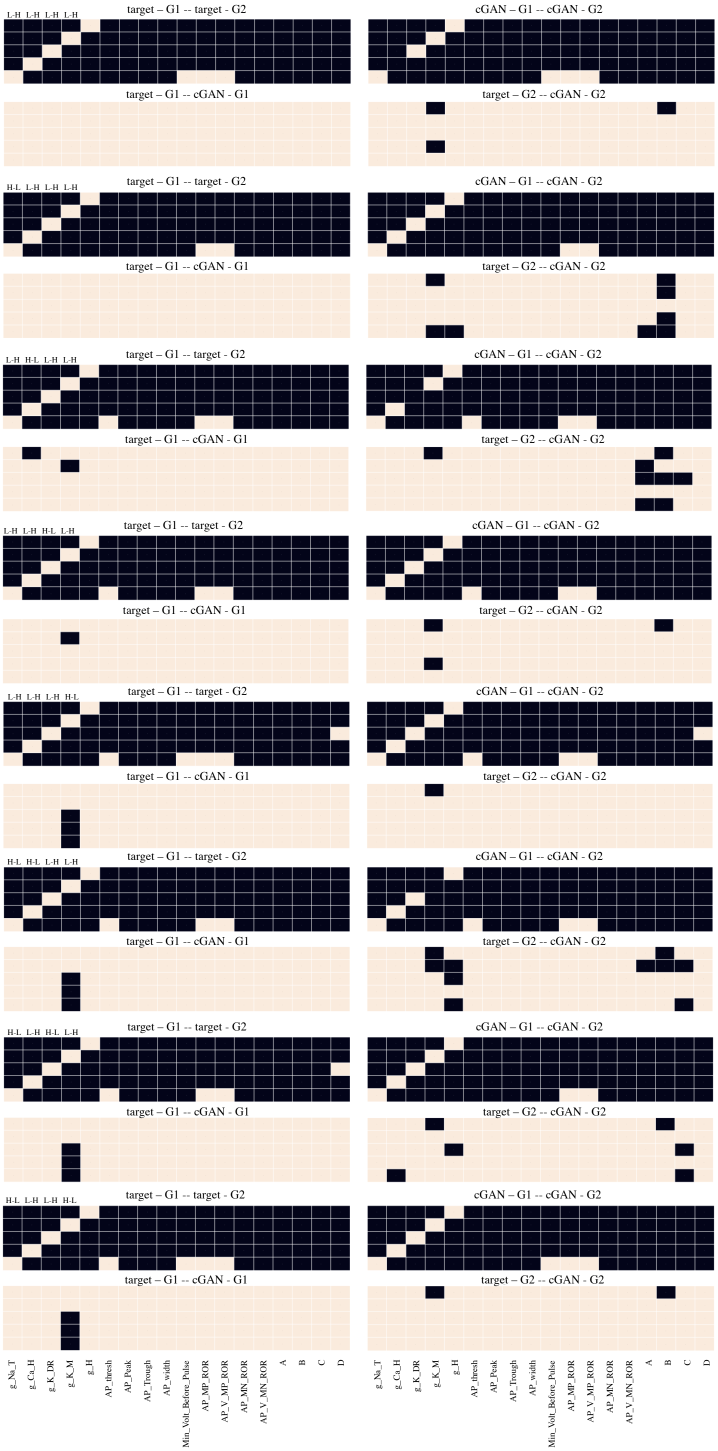


**Fig. S5 5 choose 4 - KS tests.** Panels are arranged in a similar fashion as Fig. S2. From top to bottom: *(1) **L-H L-H L-H L-H**, (2) **H-L L-H L-H L-H**, (3) **L-H H-L L-H L-H**, (4) **L-H L-H H-L L-H**, (5) **L-H L-H L-H H-L**, (6) **H-L H-L L-H L-H**, (7) **H-L L-H H-L L-H**, (8) **H-L L-H L-H H-L**.*

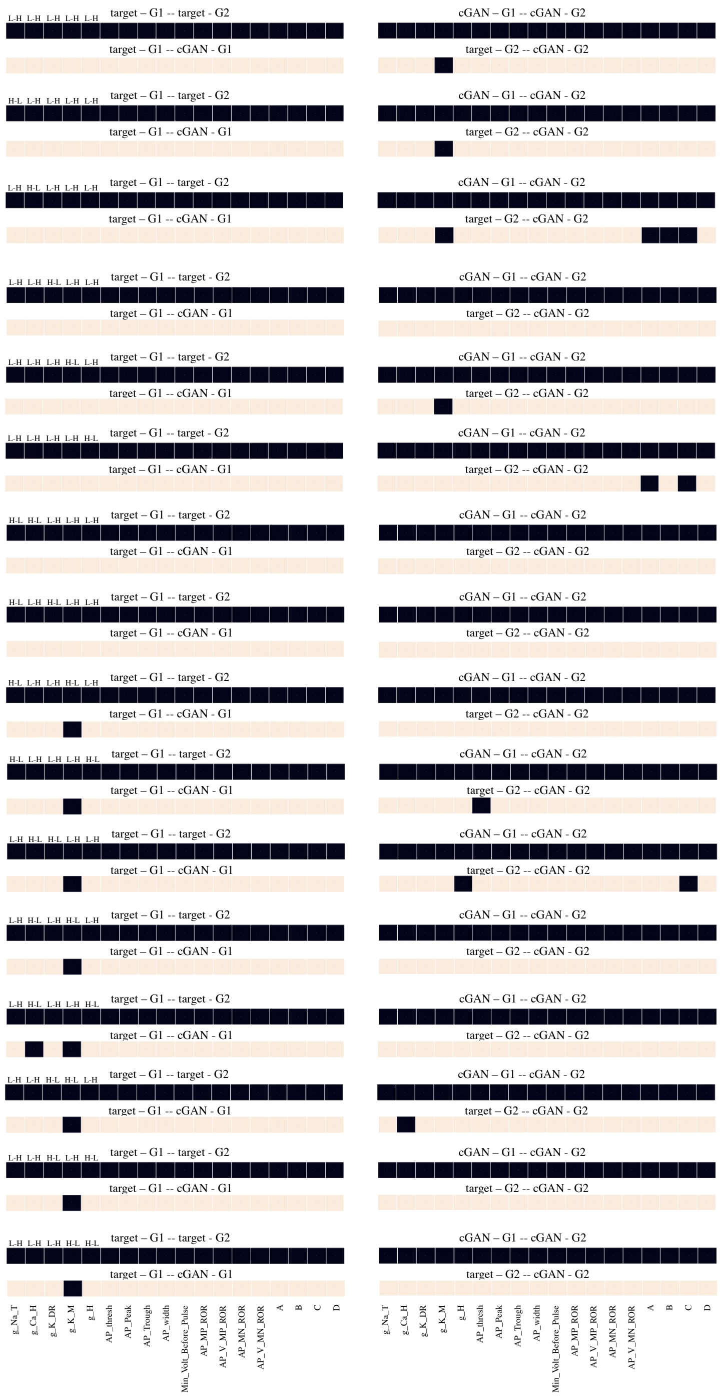

**Fig. S6 5 choose 5 - KS tests.** - Panels are arranged in a similar fashion as Fig. S2. From top to bottom: *(1) L-H L-H L-H L-H L-H*, *(2) H-L L-H L-H L-H L-H*, *(3) L-H H-L L-H L-H L-H*, *(4) L-H L-H H-L L-H L-H*, *(5) L-H L-H L-H H-L L-H*, *(6) L-H L-H L-H H-L L-H H-L*, *7) H-L H-L L-H L-H L-H*, *(8) H-L L-H H-L L-H L-H*, *(9) H-L L-H L-H H-L L-H*, *(10) H-L L-H L-H L-H H-L*, *(11) L-H H-L H-L L-H L-H*, *(12) L-H H-L L-H H-L L-H*, *(13) L-H H-L L-H L-H H-L*, *(14) L-H L-H H-L H-L L-H*, *(15) L-H L-H H-L L-H H-L*, *(16) L-H L-H L-H H-L H-L*.

5-choose-1

| | |
|---|---|
| True | 2/90 |
| 0/90 | 3/90 |

Total = 5/270

5-choose-2

| | |
|---|---|
| True | 7/360 |
| 3/360 | 15/360 |

Total = 25/1080

5-choose-3

| | |
|---|---|
| True | 9/720 |
| 9/720 | 38/720 |

Total = 56/2160

5-choose-4

| | |
|---|---|
| True | 7/720 |
| 15/720 | 41/720 |

Total = 63/2160

5-choose-5

| | |
|---|---|
| True | 0/288 |
| 9/288 | 13/288 |

Total = 22/864

**Fig. S7 Summary of KS test results for all 5 choose $k$ synthetic target data cases.** The demoninators are the total number of tests, and the numerators are the number of those tests for which the null hypothesis was rejected. **Top left quadrants:** These KS tests are taken to be the ground truth as they compared target G1 versus target G2. **Top right quadrants:** These KS tests compared cGAN-G1 versus cGAN-G2. **Bottom left quadrants:** These KS tests compared target-G1 versus cGAN-G1. **Bottom left quadrants:** These KS tests compared target-G2 versus cGAN-G2.

#### 3.3 Output of cGAN samples in feature space for synthetic target data

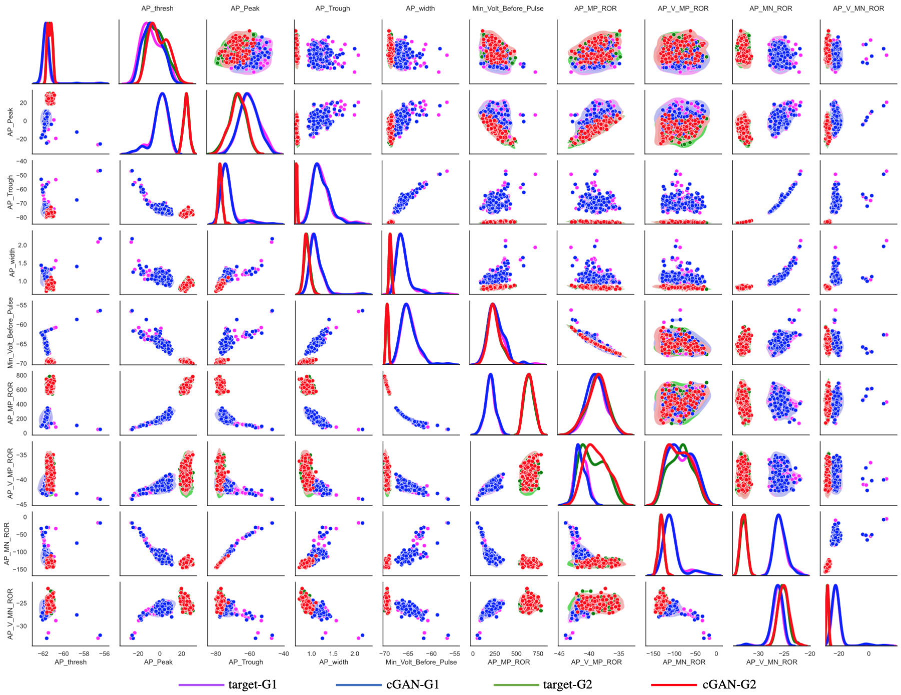


**Fig. S8 Performance of cGAN on synthetic targets from 2 groups with distinct parameter structures - AP features.** KDE plots (main diagonals) and scatter plots (lower and upper triangles) for Group 1 (G1) target data (magenta), Group 2 (G2) target data (green), cGAN samples for G1 (blue) and cGAN samples for G2 (red). *Lower main diagonal and lower triangle* - only 1 parameter ($g_{NaT}$) is distributed differently in the G1 target data than in the G2 target data, and the other 4 parameters have the same distribution in the G1 and G2 target data. We refer to this scenario as "5 choose 1" in the Supplementary Methods. *Upper main diagonal and upper triangle* - Four parameters (all parameters except $g_{NaT}$) are distributed differently in the G1 target data than in the G2 target data. We refer to this scenario as "5 choose 4" in the Supplementary Methods.

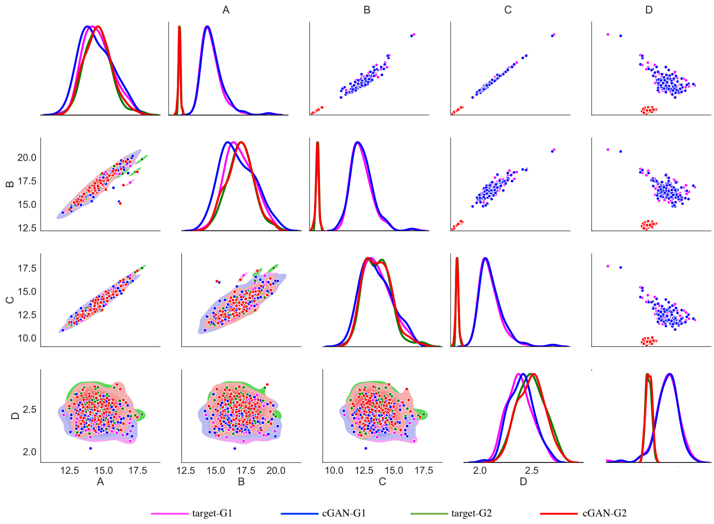


**Fig. S9 Performance of cGAN on synthetic targets from 2 groups with distinct parameter structures - HP features.** KDE plots (main diagonals) and scatter plots (lower and upper triangles) for Group 1 (G1) target data (magenta), Group 2 (G2) target data (green), cGAN samples for G1 (blue) and cGAN samples for G2 (red). *Lower main diagonal and lower triangle* - only 1 parameter ($g_{NaT}$) is distributed differently in the G1 target data than in the G2 target data, and the other 4 parameters have the same distribution in the G1 and G2 target data. *Upper main diagonal and upper triangle* - Four parameters (all parameters except $g_{NaT}$) are distributed differently in the G1 target data than in the G2 target data.

#### 3.4 Output of cGAN samples in feature space for experimental target data

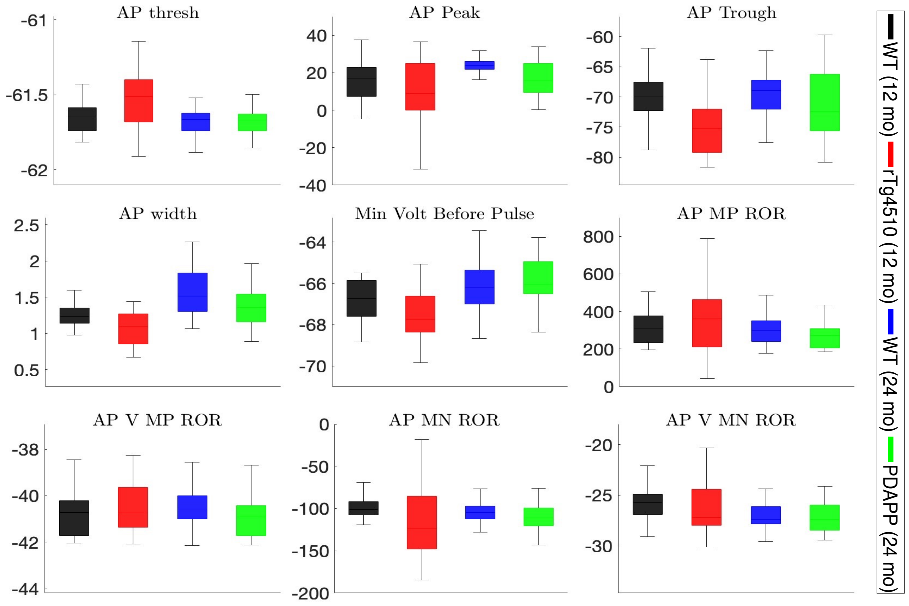


**Fig. S10** Box and whisker plots of the action potential (AP) features extracted from the mechanistic model voltage traces obtained by pushing forward the cGAN parameter samples with experimental target data.

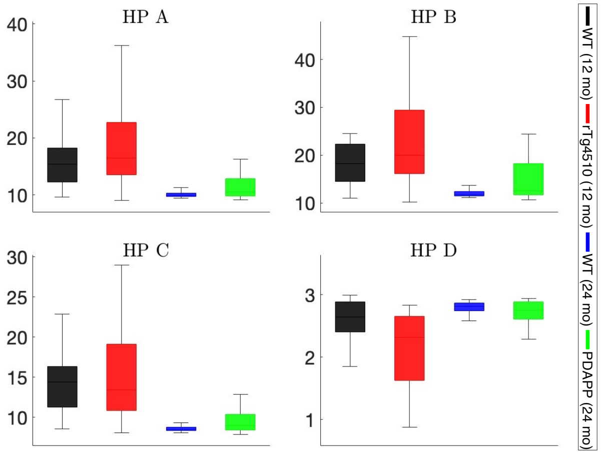


**Fig. S11** Box and whisker plots of the membrane hyperpolarization (HP) features extracted from the mechanistic model voltage traces obtained by pushing forward the cGAN parameter samples with experimental target data.
